# Balancing selection and the functional effects of shared polymorphism in cryptic *Daphnia* species

**DOI:** 10.1101/2024.04.16.589693

**Authors:** Connor S. Murray, Madison Karram, David J. Bass, Madison Doceti, Dörthe Becker, Joaquin C. B. Nunez, Aakrosh Ratan, Alan O. Bergland

**Affiliations:** Department of Biology, University of Virginia, Charlottesville, VA, USA; School of Biosciences, Ecology and Evolutionary Biology, University of Sheffield, Sheffield, UK; Center of Public Health Genomics, University of Virginia, Charlottesville, VA, USA; Department of Public Health Sciences, University of Virginia, Charlottesville, VA, USA

**Keywords:** Shared polymorphism, *Daphnia*, balancing selection, incomplete lineage sorting, hybridization, convergent evolution, overdominance, opsins

## Abstract

The patterns of genetic variation within and between related taxa represent the genetic history of a species. Shared polymorphisms, loci with identical alleles across species, are of unique interest as they may represent cases of ancient selection maintaining functional variation post-speciation. In this study, we investigate the abundance of shared polymorphism in the *Daphnia pulex* species complex. We test whether shared mutations are consistent with the action of balancing selection or alternative hypotheses such as hybridization, incomplete lineage sorting, or convergent evolution. We analyzed over 2,000 genomes from North American and European *D. pulex* and several outgroup species to examine the prevalence and distribution of shared alleles between the focal species pair, North American and European *D. pulex*. We show that while North American and European *D. pulex* diverged over ten million years ago, they retained tens of thousands of shared alleles. We found that the number of shared polymorphisms between North American and European *D. pulex* cannot be explained by hybridization or incomplete lineage sorting alone. Instead, we show that most shared polymorphisms could be the product of convergent evolution, that a limited number appear to be old trans-specific polymorphisms, and that balancing selection is affecting young and ancient mutations alike. Finally, we provide evidence that a blue wavelength opsin gene with trans-specific polymorphisms has functional effects on behavior and fitness in the wild. Ultimately, our findings provide insights into the genetic basis of adaptation and the maintenance of genetic diversity between species.

## Introduction

Genetic diversity reflects a species’ history and serves as the foundation for adaptation to ecological change. In nature, mutations arise, and their persistence time is a function of their selective value and the effective population size of the focal species (Crow & Kimura, 1970). One distinct type of genetic variant is a shared polymorphism, in which mutations are identical by state across closely related species (Wang & Mitchell-Olds, 2017). The abundance and frequency of shared polymorphisms between two species can provide insight into some of the most interesting processes in evolution. Shared polymorphism that arose prior to the split of two species are generally referred to as trans-species polymorphism (Hedrick, 2013; Wiuf et al., 2004; Wu et al., 2017). Trans-specific polymorphisms can be used to study the speciation process (Klein et al., 1998), helping refine estimates of the timing and population sizes at divergence (Edwards et al., 2000). Unless divergence happened recently or there is ongoing gene flow, it is unlikely that neutral polymorphism will be retained in both species for long (Leffler et al., 2013). Therefore, the presence of shared polymorphism between two species with limited gene-flow can be a powerful way to identify balanced polymorphisms (Clark, 1997). These polymorphisms are presumed to be maintained by temporal or spatial variation in the direction of natural selection (Bergland et al., 2014; Ségurel et al., 2012, 2013), or by genetic overdominance (Wills, 1975). Shared polymorphisms can also indicate convergent adaptive evolution (Castoe et al., 2009) or adaptive introgression (Hedrick, 2013), and these polymorphisms themselves can also be the target of balancing selection (Wang & Mitchell-Olds, 2017).

Identifying the forces that generate and maintain shared polymorphism is therefore an important problem in evolutionary genetics. However, identifying the contribution of neutral, demographic, and adaptive evolutionary processes to the generation and maintenance of shared polymorphisms is challenging. This is especially so for young species because there has not been enough time for species-specific alleles to fix, and for drift to erode shared polymorphism present due to incomplete lineage sorting. In contrast, testing alternative hypotheses for the generation of shared polymorphism can sometimes be more tractable for slightly older species pairs. This is because neutral trans-species polymorphisms are expected to be rare thereby eliminating incomplete lineage sorting as a main driver of shared polymorphism. If sufficient time has occurred for fixation of species-specific alleles, then adaptive introgression will be relatively easy to identify (Huerta-Sánchez et al., 2014) especially if it occurred recently. However, if only a single trans-specific polymorphism is functional, recombination will erode the ancestral haplotypes (Gao et al., 2015) and gene trees will align with species trees causing ambiguity when differentiating between convergent evolution and trans-specificity (Unckless et al., 2016). In contrast, long-term balancing selection of a trans-specific polymorphism can be relatively unambiguous if there are multiple sites at a locus that are shared polymorphisms, and these are tightly linked causing allele trees to not align with species trees (Wang et al., 2020). The presence of trans-specific haplotypes suggests that multiple functional sites at the locus are the target of some form of balancing selection (Charlesworth, 2006).

*Daphnia* species are an excellent model to study the mechanisms that generate and maintain shared polymorphism. *Daphnia* are freshwater microcrustaceans that have been the focus of ecological and evolutionary research for over a century (Ebert, 2022). Among the most widely studied taxa within this genus are *D. magna* (Decaestecker et al., 2007), *D. obtusa* (Spitze, 1993), as well as *D. pulex* (Lynch et al., 2017) and its close relatives (Colbourne et al., 2011). The *D. pulex* species group is currently in the process of an adaptive radiation (Fryer, 1991). Owing to their recent divergence time, some members of the North American *D. pulex* species group are known to hybridize in the wild (Held et al., 2016), resulting in contagious obligate asexuality (Xu et al., 2015). Members of the *D. pulex* species group, including *D. obtusa*, are found across the Palearctic and Nearctic (Crease et al., 2012), and recently established populations can be found in other regions of the world (So et al., 2015). Although *D. obtusa*, *D. pulicaria*, and *D. pulex* have been identified on multiple continents, each of these three taxa represent polyphyletic groups (Černý & Hebert, 1999). For instance, based on mitochondrial sequence, *D. pulex* found in North America is more closely related to North American *D. pulicaria* than it is to European *D. pulex* (Crease et al., 2012). The confusion of species identification and naming in this genus is due to the similar morphology (Dodson, 1981) and ecological niches of these taxa (Chin & Cristescu, 2021) plus their capacity to interbreed (Pantel et al., 2011), generally reflecting the taxonomic ambiguities within the species group (Hebert & Wilson, 1994), and among zooplankton in general (Brooks, 1957).

The evolutionary and ecological history of the *D. pulex* species group affords us an ideal opportunity to study the evolutionary forces that have shaped patterns of shared polymorphism. Here, we assessed alternative evolutionary mechanisms that can generate and maintain shared polymorphisms between North American and European *D. pulex*. Using population genomic data from samples in North America and Europe, we first confirm that North American and European *D. pulex* are distinct species that have diverged millions of years ago. Next, we show that North American and European *D. pulex* possess tens of thousands of shared polymorphisms, whose abundance cannot be explained by incomplete lineage sorting, hybridization, introgression, or gene-flow. Therefore, we conclude that many of these shared polymorphisms arose either via convergent evolution or have been maintained since the split between these taxa. For many shared polymorphisms, we cannot differentiate which of these two mechanisms is most likely. However, a limited number of genes show a strong excess of shared polymorphisms that are in linkage disequilibrium, consistent with long-term balancing selection operating on a haplotype. One of these genes is a single-copy blue wavelength opsin, part of a gene family that has previously been identified as a target of rapid adaptive evolution in *Daphnia* (Brandon et al., 2017; Ye et al., 2023). We show that European *D. pulex* clones harboring alternate genotypes for this blue wavelength opsin have differences in movement and activity that is dependent on light conditions and provide evidence for overdominance in the wild. Taken together, our results highlight the abundance, selective history, and function of shared polymorphisms in *Daphnia* and contributes to the understanding of the phylogeography for this classic model system.

## Materials and Methods

### Sampling and sequencing European Daphnia genomes

*Daphnia* were sampled from 16 ponds throughout England in 2018. Samples were transported to the University of Virginia and clonally derived isofemale lines were established. Samples were identified as either *D. pulex*, *D. pulicaria*, or *D. obtusa* based on morphological characteristics using an online dichotomous key (http://cfb.unh.edu/cfbkey/html/anatomy/daphnia/daphnia.html). DNA extraction and library preparation followed methods outlined in Kubow *et al*. (2022). Briefly, for each isofemale line multiple individuals were exposed to antibiotics (streptomycin, tetracycline, and ampicillin, 50 mg/L of each) and fed Sephadex G-25 beads to clear their gut of algae. Samples were homogenized using metal beads and a bead beater and DNA was extracted using the Agencourt DNAdvance kit (Beckman-Coulter). RNA was removed using RNase followed by an additional bead cleanup. DNA was quantified using the broad-range Quant-iT dsDNA kit (ThermoFisher Scientific) and normalized to 1 or 2 ng/μL before library construction. Full genome libraries were constructed using a scaled down Nextera protocol (Baym et al., 2015). Libraries were size selected for fragments ranging from 450 to 550 bp using a Blue Pippin and quality checked using a BioAnalyzer. Samples were sequenced on a HiSeq X platform, paired-end 150bp.

### Publicly available Daphnia genomes

Genome sequences of North American and European *D. pulex*, *D. pulicaria*, and *D. obtusa* were obtained from NCBI’s Sequence Read Archive (SRA; Leinonen et al., 2011). We incorporated wild-sequenced or isogenic female lineages (Barnard-Kubow et al., 2022; Lynch et al., 2017; Xu et al., 2015; Ye et al., 2022), and excluded samples that were from mutation accumulation studies. Species identity for these samples was based on annotations provided in each SRA record.

### Short-read mapping

Prior to short-read mapping of all samples, sequencing adaptors were removed using *trimmomatic v0.39* (Bolger et al., 2014), and overlapping reads were merged using *pear v0.9.11* (Zhang et al., 2014). All samples were mapped to the European *D. pulex* genome (Barnard-Kubow et al., 2022) using *bwa mem v0.7.17* (H. Li, 2013), and downstream data manipulation was performed using *samtools merge v1.12* (H. Li et al., 2009). Duplicate reads for every bam file were marked and removed using *picard v2.23.4* (https://broadinstitute.github.io/picard/). Quality control metrics were assembled using *fastqc v0.11.5* (Andrews, 2010) and *MultiQC v1.11* (Ewels et al., 2016). S*amtools flagstat* counted the mapped and properly paired reads.

Additionally, we mapped North American *D. pulex* to the North American *D. pulex* reference genome (KAP4; GenBank assembly: GCF_021134715.1) using the same mapping strategy outlined above. We created a liftOver file to translate features in KAP4 to D84A. The chains exhibited good coverage, allowing us to translate 72.6% of the KAP4 genome to D84A (Lee et al., 2022). We created the liftOver file by running pairwise alignments using *lastz* followed by the use of various UCSC tools to chain the alignments, sort them, filter them, and convert them into UCSC nets and chains (Harris, 2007). We used *LiftOverVCF* from *picard* to convert the KAP4 aligned VCF to the D84A genome coordinates. We then assessed the concordance of the SNP classifications between North American samples mapped to D84A and the liftOver VCF (Supplemental Figure 1C-D).

All analyses using North American and European *D. pulex, D. pulicaria,* and *D. obtusa* were conducted using samples mapped to the European *D. pulex* reference genome. We assessed reference allele bias for these interspecific mappings by calculating the proportion of alternative and reference allele dosage for 1,000 biallelic heterozygous BUSCO gene SNPs across the genome (with 100 bootstrap resampling). All analyses focusing exclusively on shared polymorphisms between European and North American *D. pulex* were conducted using the intersection of SNPs identified by mapping North American *D. pulex* and European *D. pulex* to their respective reference genomes.

### SNP calling and filtering

We used *HaplotypeCaller* and *GenotypeGVCFs* from *gatk v4.1.6.0* to create VCF files (Poplin et al., 2017). *VariantFiltration* in *gatk* removed low-quality SNPs recommended for organisms without reference panels: (“QD<2.0”, “QUAL<30.0”, “SOR>3.0”, “FS>60.0”, “MQ<40.0”, “MQRankSum<-12.5”, “ReadPosRankSum<-8.0”). We removed sites flanking ±10bp any indels using *bcftools filter --SnpGap 10* (Li et al., 2009) and removed indels using *SelectVariants* in *gatk*. We annotated SNPs using *snpEff v4.3t* (Cingolani et al., 2012).

Samples with average genome-wide missingness >10% were removed from analyses as was any genomic region with more than 10% missingness across the remaining samples. We removed regions with high (DP≥35) and low mean site read depth (DP≤8), along with chromosomal endpoints, regions of the reference genome with large stretches of gaps, and regions of Ns as described in Barnard-Kubow et al., 2022. Repetitive elements identified in the European *D. pulex* genomes were classified with *RepeatMasker v4.0.8* and were removed (Tarailo-Graovac & Chen, 2009). We restricted analyses to the genic and non-genic regions associated with the 6,544 single-copy ortholog genes between European and North American *D. pulex* from *OrthoFinder v5* (Emms & Kelly, 2019) for the SNPs that were retained from the liftOver. Most analyses removed SNPs that have a minor allele frequency (MAF) less than 0.01 within-species. After filtering, 347,200 SNPs represent the whole-genome SNP set. We restricted phylogenetic analyses to BUSCO genes identified with *Panther* annotations (Mi et al., 2013; Seppey et al., 2019; Simão et al., 2015). The BUSCO gene SNP set includes 138,024 SNPs. Principal component analysis (PCA) of SNPs was conducted in *SNPRelate v1.24.0* while excluding sites with MAF<0.01 (Zheng et al., 2012). D_xy_ was calculated using *PopGenome v2.7.5* (Revell, 2019).

### Assigning multi-locus genotypes

Every sample was assigned to a multi-locus genotype (MLG) using the *poppr v2.9.3* package (Kamvar et al., 2014) in *R v4.0.3*. Unless otherwise noted, every analysis was subset based on picking a representative sample with the highest coverage for each MLG (Supplemental Table 1).

### Mitochondrial tree

We annotated the D84A mitochondrion using *MITOs v1* (Bernt et al., 2013). We aligned and called SNPs using *bcftools mpileup v1.9* and *bcftools call*. We excluded reads that had low-quality scores (Q<20) and high depth (DP>100) using *bcftools filter*. And generated consensus FASTA files using *bcftools consensus*. We mapped North American *D. pulex* and *D. pulicaria* to the North American *D. pulex* mitochondrial genome sequence (GenBank accession: NC_000844.1) and mapped both North American and European *D. obtusa* samples to the North American *D. obtusa* mitochondrial genome sequence (GenBank accession: CM028013.1). The mitochondrial sequence of European *D. magna* was used as an outgroup (GenBank accession: NC_026914.1). We assembled homology blocks using *exonerate v2.4.0* (Slater & Birney, 2005) for the 13 protein-coding genes and found high sequence similarity (>80%), except for *atp8*. Therefore, we assembled trees excluding *atp8*. We then used *mafft v7.475* (Katoh & Standley, 2013) to assemble multiple sequence alignments (MSA). We concatenated these MSAs for each gene using *seqkit concat v2.2.0* (Shen et al., 2016) and ran *iqtree2 v2.1.2* with 1,000 bootstraps (Supplemental Figure 2; Hoang et al., 2018; Kalyaanamoorthy et al., 2017).

### Estimating divergence-time

We used *Snapp v1.6.1* within *Beast2 v2.6.6* to estimate the split-time within the species complex (Bouckaert et al., 2014). We used two representative individuals with the highest coverage for each species. We used 3,000 randomly sampled BUSCO gene SNPs, 1 million iterations, and a 10% burn-in. The output tree was time-constrained for the outgroup species, European *D. obtusa,* to 31 million years ago (MYA with a confidence interval of 1 MYA based upon a genus-wide tree; Chin & Cristescu, 2021; Cornetti et al., 2019). We used *Tracer v1.7.1* to quantify MCMC convergence (Rambaut et al., 2018).

### Hybridization statistics

We used *ADMIXTURE v1.3.0* (Alexander & Lange, 2011), excluding any sites with MAF<0.01 and thinned every 500 SNPs. We varied the number of clusters (*k*) from 2-25 and calculated the cross-validation error (CV) at every *k* model. We chose *k*=9 because the minimum CV score was reached. We quantified the magnitude of introgression using *Dsuite v0.5* (Malinsky et al., 2021) with European *D. obtusa* as the outgroup.

### Historic N_e_ and demographic inference of migration

To calculate historical *N_e_* for European and North American *D. pulex,* we ran *MSMC2 v2.1.1* and *SMC++ v1.15.4* (Schiffels & Wang, 2020; Terhorst et al., 2017). We performed demographic inference with *moments v1.1.0* in *python3* (Jouganous et al., 2017). We tested two models: one with-migration and one without-migration. For the former model, we used *moments*’ *split_mig* model. For the latter, we used *split_mig* with no migration. We ran inference on 20x20 SFS projections until model convergence and classified a shared polymorphism as an allele whose allele frequency is above 1/20 in both species. We used a mutation rate *μ*=5.69x10^-9^ (Lynch et al., 2017). We followed the methods of McCoy et al., (2014) to convert coalescent units of into standard units. We estimated the ancestral population size as *N_ANC_*=200,000/*η_EU_*, where *η_EU_*=*N_e_* European *D. pulex* is an approximate from historic demographic inference (Supplemental Figure 3 left panel). Then, with *moments*’ estimates *η_NA_* for *N_NA_*, τ for *t_split_*, and *M* for migration, we calculated: *N_NA_*=*N_ANC_*×*η_NA_*, *N_EU_*=*N_ANC_*×*η_EU_*, *t_split_*=2*N_ANC_*×*τ*, and *m*=*M÷*2*N_ANC_*. We chose to associate European *D. pulex* with the ancestral species because the reference genome isolate is a European *D. pulex* clone.

### Classifying shared polymorphisms between North American and European D. pulex

We classified each mutation as a fixed difference between species, polymorphic within-species, or a shared polymorphism between species. We classified sites as polymorphic (within species or shared) if the minor allele frequency in either or both species was greater than 0.01 (Supplemental Table 2).

We tested whether the extent of shared polymorphism can be explained by incomplete lineage sorting using methods outlined elsewhere (Novikova et al., 2016; Wiuf et al., 2004). The formula in Novikova et al., (2016) estimates the number of expected shared polymorphisms between species, where *d*_between_ is D_xy_ between North American and European *D. pulex*, and *d*_NAm_ & *d*_Euro_ are within-species polymorphism.

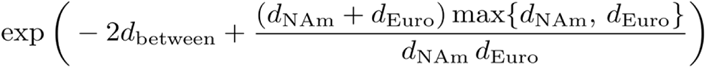

### Balancing selection statistics

*BetaScan v1* was used to calculate *β^1^* statistic within species using the folded site frequency spectrum (Siewert & Voight, 2017). The *α_b_* statistic tested for the proportion of sites under balancing selection between species-pairs from Soni et al., 2022. Where *Poly.* are SNPs within-species and *SP* are shared polymorphisms between-species. *SYN* are synonymous sites and *NS* are non-synonymous sites:

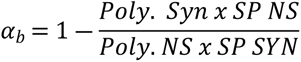

### Phylogenetic tree test using pairwise cophenetic distances

We tested the local sequence genealogy to test for trans-specificity versus convergent evolution (Koenig et al., 2019; Nunez et al., 2021). This test used trees built from 500bps flanking high-frequency non-synonymous shared polymorphisms (MAF>0.1). We calculated the median pairwise cophenetic distances (CPD) between samples (Cardona et al., 2013). We extracted haplotypes from a *WhatsHap v1.1* phased VCF (Martin et al., 2023) from 30 high-read depth individuals for each species. We chose 30 samples to keep the sample size consistent across species while decreasing model convergence time. We aligned the parental haplotypes (n=60 per species) using *mafft* and built trees using *iqtree2* (1,000 bootstraps). The null hypothesis was that the tree would be concordant with the species-tree topology. The alternative was that the tree would be discordant with the species-tree and that median CPD between North American and European *D. pulex* is higher within-species than between-species (i.e., CPD_Within_>CPD_Between_). CPD_Within-Between_=CPD_Within_-CPD_Between_, where CPD_Within_=within-species, CPD_Between_=between-species. Positive CPD_Within-Between_ indicates an allele-specific topology and negative CPD_Within-Between_ indicates a concordant topology with the species-tree. A cartoon depicting these hypotheses is in Figure 4A.

### Light exposure experiments on Daphnia activity

We developed a behavioral assay to collect activity data on 12 distinct European *D. pulex* clones using a DAM Trikinetics monitor (Chiu et al., 2010). In total, we measured activity for 216 individuals. The Trikinetics monitor has 32 wells filled with 5mm diameter plastic tubes. Each well has an infrared light beam that when broken by a *Daphnia* individual will count as an activity event. We exposed individuals to white light, blue light, and dark lighting conditions using blackout boxes mounted with LEDs (described in Erickson et al., 2020). Individual *Daphnia* were placed inside a plastic tube with artificial pond water media (ASTM; Standard, 2007) while each Trikinetics monitor collected activity measurements over a twelve-hour experimental period, sampling every 5 seconds. We excluded measurements during the first hour to allow individuals to settle in. For 95% of the 5-second intervals, 0 or 1 beam break was recorded and 99.9% of intervals had 4 or fewer beam breaks. Therefore, for each 5-second interval, we converted the number of beam breaks recorded into a binary variable (>=1 beam-break vs 0 beam-breaks) and calculated total activity as the fraction of 5-second intervals with more than one beam break per individual over the course of the experiment. We modeled total activity with a generalized linear mixed effect model using *lme4 v1.1-27.1* in *R* (Bates et al., 2015) and performed likelihood ratio tests between the following models:

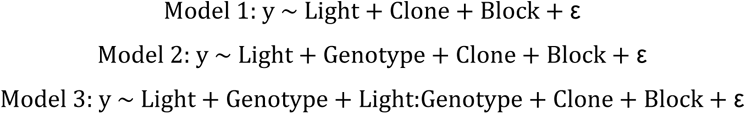

Where *y* is the fraction of intervals with activity, *Light* is the fixed effect of light treatment (white, blue, dark), *Genotype* is the fixed effect of genotype at the blue wavelength opsin (BLOP) locus, *Light:Genotype* is the fixed interaction effect, (1|*Clone*) is the random effect of clone, (1|*Block*) is the random effect of one of the three experimental blocks run over successive weeks, and ε is the binomially distributed error with weights equal to the number of 5-second intervals (ca. 7800). We conducted likelihood ratio tests between Model 1, Model 2, and Model 3 using the *anova()* function in *R* (Supplemental Table 3). In addition, we performed an additional analysis that explicitly models elapsed time in the experiment as a fixed effect and includes the individual *Daphnia* identifier as a random effect to account for repeated measures. The results of that analysis are in line with the more straightforward model presented here and we show those results in Supplemental Table 4.

### Daphnia11806-RA orthologs

We tested the orthology of *Daphnia11806-RA* by BLASTing the amino acid sequence against the NCBI database using *blastp v2.13.0* (Sayers et al., 2022).

### Statistics and visualization

Analyses were performed using *R v3.6.2–4.0.3* (R Core Development Team 2013). We used the following packages for analysis and visualization: *tidyverse v1.3.1* (Wickham et al., 2019), *ggplot2 v3.3.5* (Villanueva & Chen, 2019), *ggtree v2.0.4* (Xu et al., 2022), *ape v5.4-1* (Paradis & Schliep, 2019), *patchwork v1.0.1* (Thomas Lin Pedersen, 2022), *viridis v0.5.1* (Garnier et al., 2021), *data.table v1.12.8* (Dowle & Srinivasan, 2023), *foreach v1.4.7*, *doMC v1.3.5* (Daniel et al., 2022), *SeqArray v1.26.2* (Zheng et al., 2017).

### Data availability

The D84A mitochondrion was uploaded to NCBI (JAHCQT000000000) and updated to the existing accession: GCA_023526725.1. The novel 93 genomes described here were uploaded to NCBI under the accession: PRJNA982532. The metadata for samples is located in Supplemental Table 1. The VCF and GDS are deposited on dryad: https://doi.org/10.5061/dryad.dncjsxm3p. Scripts and data are deposited on GitHub: https://github.com/connor122721/SharedPolymorphismsDaphnia.

## Results

### Thousands of Daphnia genomes

We first assembled short-read genomic data for 2,321 samples of *D. pulex*, *D. pulicaria*, and *D. obtusa* collected from North American and European ponds (Figure 1A). This includes whole genomes published elsewhere (Barnard-Kubow et al., 2022; Lynch et al., 2017; Xu et al., 2015; Ye et al., 2022), along with 93 samples reported here for the first time. We aligned samples to the European *D. pulex* assembly (D84A; Barnard-Kubow et al., 2022) and identified 347,200 SNPs after filtering. In brief, our filtering methods removed regions that could prove problematic for population genomic analyses across related species. The SNP that we used represent within-species SNPs, fixed differences, and shared polymorphisms classified between North American and European *D. pulex*. Because lineages could be clonally derived from a recent common ancestor, each sample was assigned to a multi-locus genotype using the filtered SNP set (MLG; Supplemental Table 1). In all analyses, unless otherwise noted, we restricted to one sample per MLG (n=1,173).

**Figure 1.**
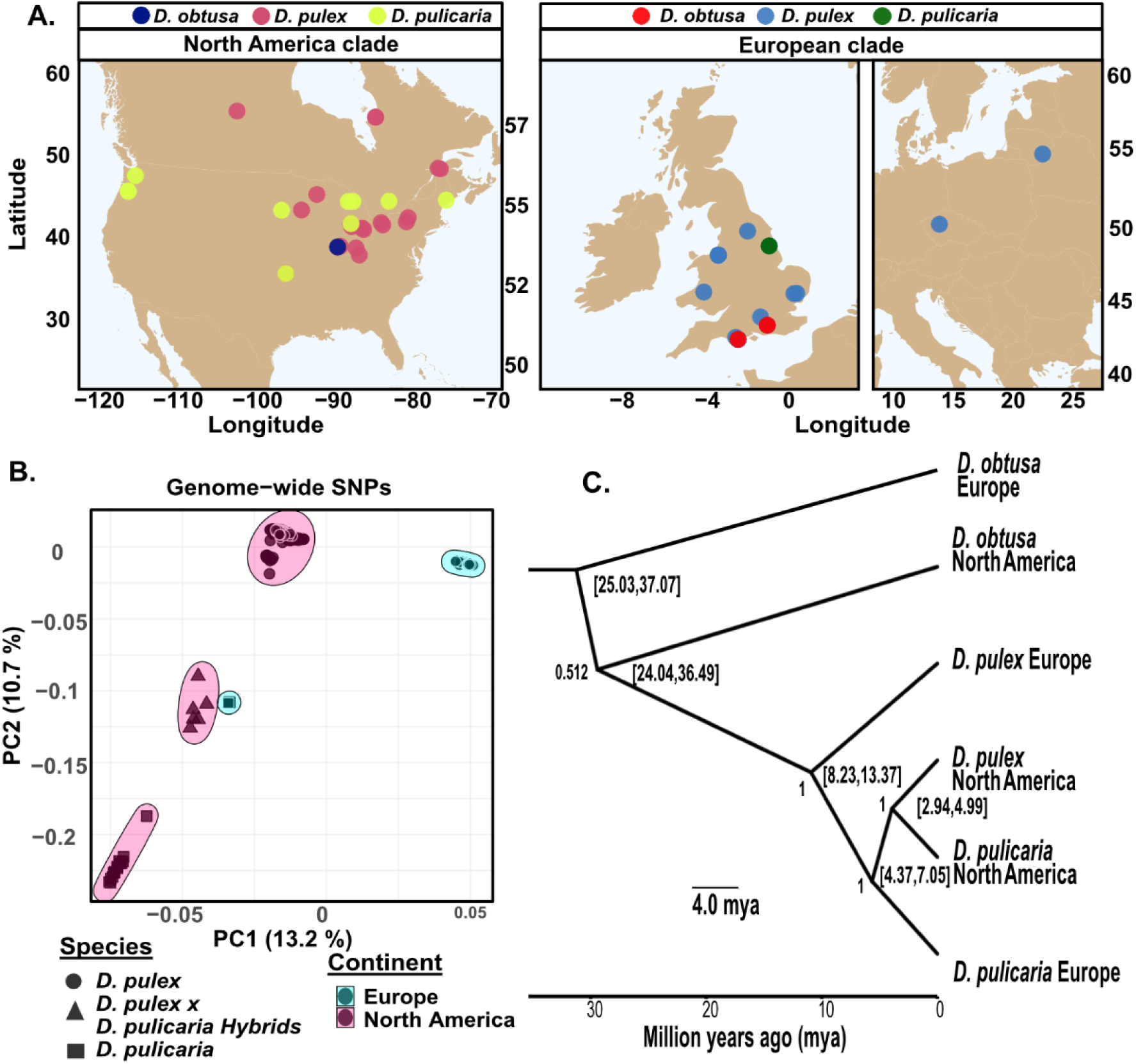
Genetic divergence of the *Daphnia pulex* species complex. **A)** Sample origin of the North American and European clades, each consisting of *D. pulex*, *D. pulicaria*, and *D. obtusa*. Most of the European clade has samples in the United Kingdom but there is one sample in both the Czech Republic and Lithuania as shown in the rightmost subfigure. **B)** The principal component axes (PC1 and PC2) using filtered genome-wide SNPs (minor allele frequency > 0.01). The proportion of variation explained by each PC is shown in parentheses. We restricted the principal component analysis to the *D.* pulex and *D. pulicaria* taxa because the *D. obtuse* taxa are so distantly related. **C)** Time-constrained phylogenetic tree restricted to 2 representative individuals within each species based on 3,000 BUSCO gene SNPs. This consensus tree is rooted with European *D. obtusa* to have 31 million years of divergence. Bracketed values are 95% confidence intervals in millions of years ago (mya). Node labels indicate the posterior probabilities estimated from 1 million bootstraps.

### Interspecific mapping does not cause systematic biases

A concern for aligning divergent sequences to the same assembly is for reference allele bias to decrease mapping efficiency and cause genotype errors (Günther & Nettelblad, 2019). To assess this, we calculated the proportion of alternative and reference allele dosage for heterozygous BUSCO gene SNPs (N=1,000; 100 bootstraps). On average, SNPs identified in North American or European *D. pulex, D. pulicaria,* or *D. obtusa* had approximately the same alternative and reference allele dosage at heterozygous sites, revealing an absence of systematic reference allele bias (Supplemental Figure 1A). Next, we mapped North American *D. pulex* samples to their species assembly (KAP4) and measured the concordance of SNP classifications between genomes. We show that 88% of SNP classifications are unchanged between assemblies (Supplemental Figure 1B&C). However, this high level of concordance could be an underestimate because of information loss incurred from lifting over assemblies (Chen et al., 2021; Günther & Nettelblad, 2019). Therefore, we conclude that the data is not systematically biased by mapping reads from non-European *D. pulex* to the European *D. pulex* assembly.

Results that highlight genetic divergence, hybridization, and introgression between taxa use the SNP classifications identified by exclusively mapping to the European *D. pulex* reference genome. To be rigorous, all results that focus on shared polymorphisms between North American and European *D. pulex* use sites that were identified as shared polymorphisms when mapping reads from each species to their respective reference genome, and then lifting over coordinates (N_SNPs_=28,983; Supplemental Table 2).

### Population genetics of the species complex

To understand the extent of divergence between species, we performed principal component analysis (PCA) on the SNP dataset after retaining sites above 0.01 minor-allele frequency (MAF) within-species (Figure 1B). The first and second PC axes are significantly different between the North American and European *D. pulex*, *D. pulicaria,* and hybrids species groups (ANOVA PC1: F_4,1154_ = 70,617, *p <* 2 x 10^-16^; ANOVA PC2: F_4,1154_ = 27,940, *p <* 2 x 10^-16^). Intriguingly, European *D. pulicaria* clusters near the known hybrids of North American *D. pulicaria* and *D. pulex* (Jackson et al., 2021; Tucker et al., 2013; Supplemental Table 1); below we test whether the samples identified as *D. pulicaria* collected in Europe are related hybrids between North American taxa or are themselves hybrids.

To evaluate the nuclear phylogeny of the *D. pulex* species complex, we built a time-constrained phylogenetic tree using BUSCO gene SNPs. The tree omitted known hybrids of North American *D. pulex* and *D. pulicaria* because they prevented model convergence. Our results show that the nodes that split the *D. pulex* species complex are generally well supported, reflecting a high pairwise sequence divergence (D_xy_) between taxa. We estimate that the split-time between North American and European *D. pulex* is around 10 million years ago (Figure 1C). The mitochondrial phylogeny also supports a reciprocally monophyletic relationship between North American and European *D. pulex*. However, North American *D. pulex* and *D. pulicaria* are not reciprocally monophyletic (Supplemental Figure 2). The recent split-time between North American *D. pulex* and *D. pulicaria* (Ye et al., 2022), their propensity to hybridize (Pantel et al., 2011), and discordant mitochondrial and nuclear phylogenies support the hypothesis that North American taxa are in the process of incipient speciation (Heier & Dudycha, 2009).

North American and European *D. pulex* possess marked differences in levels of diversity, consistent with long-term divergence. Principal component clusters are more dispersed among North American *D. pulex* than they are among European *D. pulex*, suggesting higher genetic variability within the North American clade (Figure 1B). Second, synonymous site D_xy_ between the two species is large (BUSCO genes D_xy_=0.054). Third, North American and European *D. pulex* taxa have different historic *N_e_*: the North American *D. pulex N_e_* is ∼700,000 (95% confidence intervals; 625,090.3, 772,232.5) whereas the European *D. pulex N_e_*is ∼300,000 (268,445.1, 323,248; Supplemental Figure 3).

### Hybridization in the D. pulex species group

Hybridization is common among the group of North American *D. pulicaria* and *D. pulex* species (Pantel et al., 2011), however, signals of hybridization between North American and European *Daphnia* remain less well understood. European *D. pulicaria* and North American *D. pulex*-*pulicaria* hybrids both exhibit strong signals of hybridization (Figure 2AB; *D*=0.49, *f4-ratio*=0.236, *p*=2.3x10^-16^ for European *D. pulicaria; D*=0.55, *f4-ratio*=0.48, *p*=2.3x10^-16^ for North American *D. pulex*-*pulicaria*). However, hybridization between European *D. pulicaria* and North American or closely related circumarctic species is not recent or is with other members of the complex North American *D. pulex-pulicaria* species sub-group. For example, an *ADMIXTURE* analysis reveals that European *D. pulicaria* has distinct ancestry clusters from other species, while the recent hybrids of North American *D. pulex-pulicaria* display split ancestry between North American *D. pulex* and *D. pulicaria* (Figure 2D; Alexander & Lange, 2011). We also examined heterozygosity at fixed differences between North American *D. pulex* and North American *D.* pulicaria in European *D. pulicaria* and North American *D. pulex*-*pulicaria* hybrids. These fixed differences are heterozygotes 70% of the time in North American hybrids, but only 2% of the time in European *D. pulicaria* suggesting a distinct evolutionary history of the European *D. pulicaria* clade. In summary, our findings imply that European *D. pulicaria* is likely a member of the speciose North American *Daphnia pulex* species sub-group, consistent with previous reports of a circumarctic *D. pulex* lineage predominant across Northern Eurasia (Colbourne et al., 1998).

**Figure 2.**
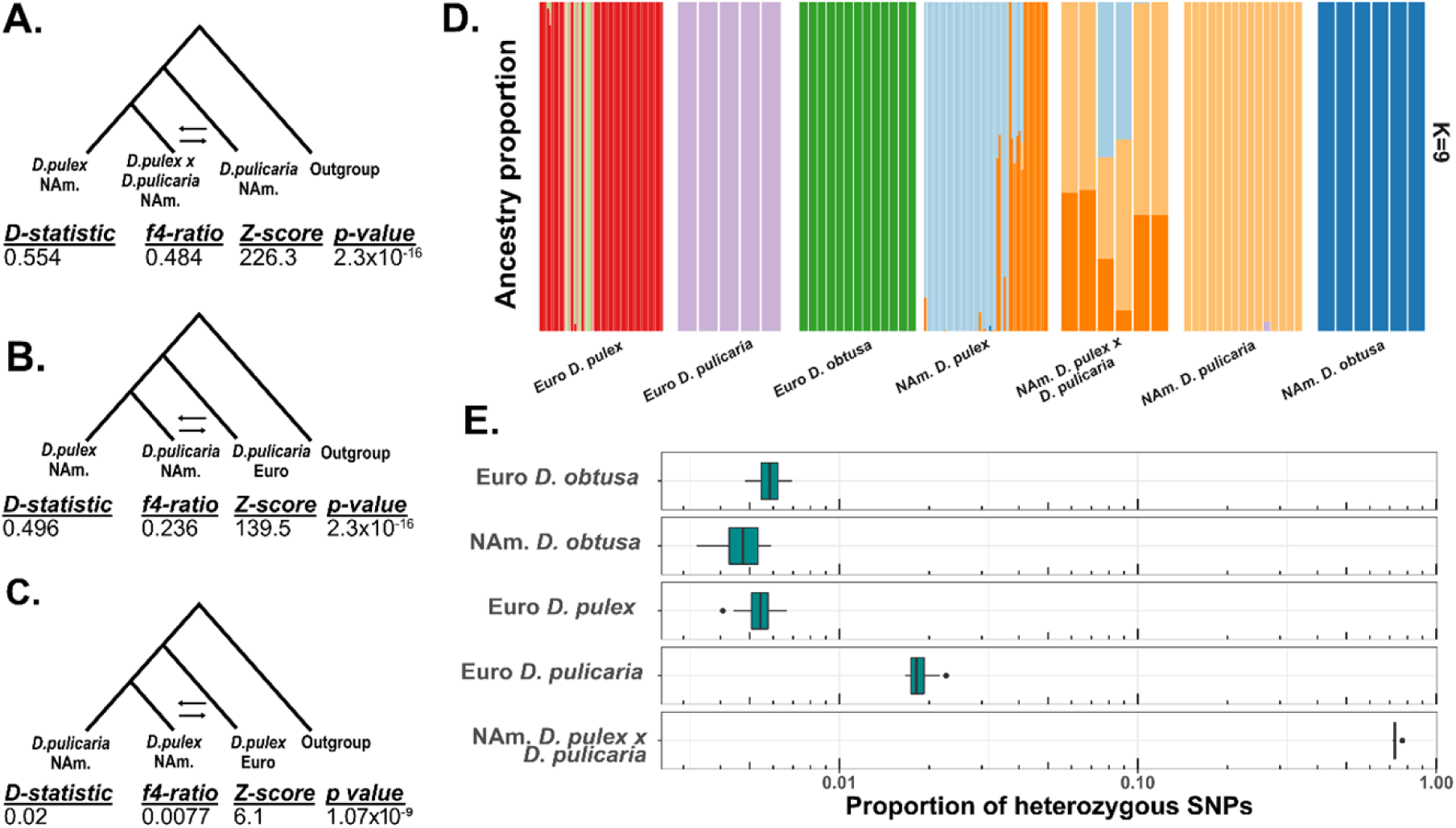
Hybridization across the *D. pulex* species complex. **A-C)** Introgressions tests using various four-species trees. The outgroup is European *D. obtusa* in all tests. *D-statistic* and *f4-ratio* describe the extent of introgression between the 2^nd^ and 3^rd^ taxa on the tree being tested. **D)** *ADMIXTURE* plot of the *D. pulex* species complex with *k*=9 having the minimal cross-validation error. Each color represents a unique ancestry group for each sample. **E)** We identified fixed differences between North American *D. pulex* and *D. pulicaria* and calculated the proportion that is heterozygous in a randomly chosen individual from the remaining taxa. The boxplot shows the distribution of these proportions from randomly sampled clones (one per MLG).

However, signals of hybridization are weak between North American and European *D. pulex* (*D*=0.02, *f4-ratio*=0.0077, *p*=1.07x10^-9^; Figure 2C). *ADMIXTURE* analysis suggests that European *D. pulex* forms several distinct ancestry groups that do not appear within any species of North American *Daphnia* (Figure 2D). Only ∼0.5% of fixed differences between North American *D. pulex* and *D. pulicaria* segregate as heterozygous sites in European *D. pulex* (Figure 2E). These results suggest that European *D. pulex* are distinct from the remaining taxa and do not have a recent history of hybridization with the other species studied.

### Extent of shared polymorphism between North American and European D. pulex is not explained by incomplete lineage sorting or migration

For species with deep split-times and low levels of migration or hybridization, we expect few shared polymorphisms to exist if such polymorphisms are neutral. For instance, based on a simple neutral model with no migration (Novikova et al., 2016; see Material & Methods) we expect to observe 336 shared polymorphisms given the split-time between North American and European *D. pulex* at synonymous sites. Yet, we observe at least 11,000 shared synonymous SNPs between these species (Supplemental Table 2).

This prediction does not account for historic migration, so we preformed demographic inference on the two-dimensional site-frequency spectrum (2D SFS) using *moments* (Figure 3A; Jouganous et al., 2017). First, we contrasted two models, one that allows constant migration (*Split + Migration*) and one where the migration rate was set to zero after population divergence (*Split*). The “*Split + Migration*” model is the best model based on the mean Bayesian information criteria (BIC) across bootstraps (“*Split + Migration*” BIC *=* 20,637, “*Split*” BIC = 33,216). Notably, the “*Split*” model severely underpredicts the number of shared polymorphisms, reflecting that incomplete lineage sorting alone is insufficient to explain the abundance of shared SNPs. The “*Split + Migration*” model itself underpredicts the number of shared polymorphisms by 25% (Figure 3A), and the model prediction shows a notable deficit of common shared SNPs and an excess of shared SNPs that are at low frequencies (Figure 3B) compared to the empirical SFS.

**Figure 3.**
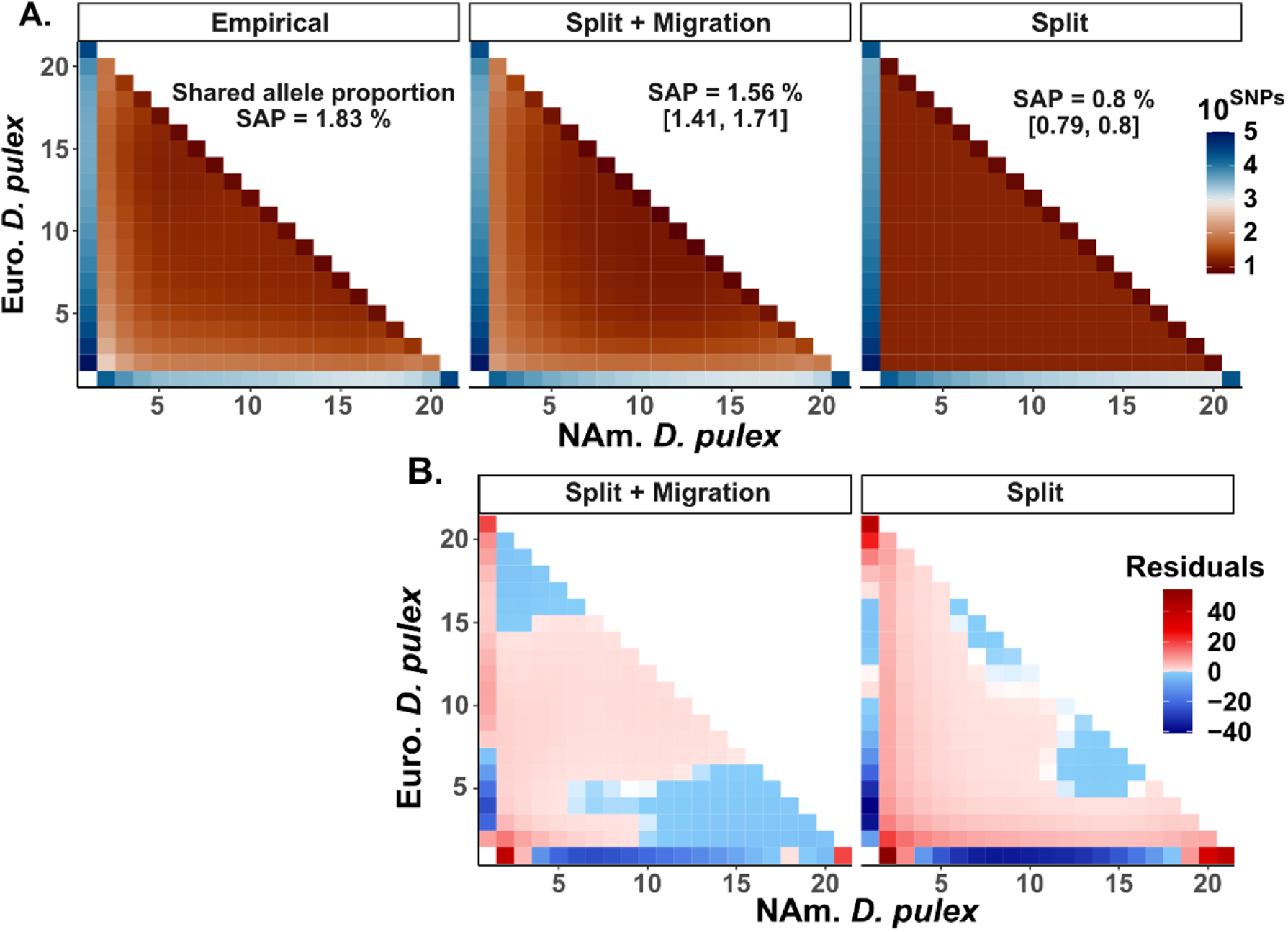
An excess of shared polymorphisms between North American and European *Daphnia pulex*. **A)** Demographic model inference between North American and European *D. pulex* based on the folded site-frequency spectrum (SFS). The empirical SFS is constructed from the genome-wide SNP dataset. The split with migration (“*Split + Migration”*) and split without migration (“*Split”*) models were generated from *moments* and we are showing the mean projection based on 1,000 bootstraps. The x and y-axis use a 20x20 SFS projection. **B)** Average standardized residuals for both models tested against the empirical SFS. Standardized residuals were calculated from the allele counts for each row and column combination of the SFS with the following formula: 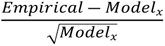, where *Model_x_* is “*Split + Migration*” or “*Split*”.

**Figure 4:**
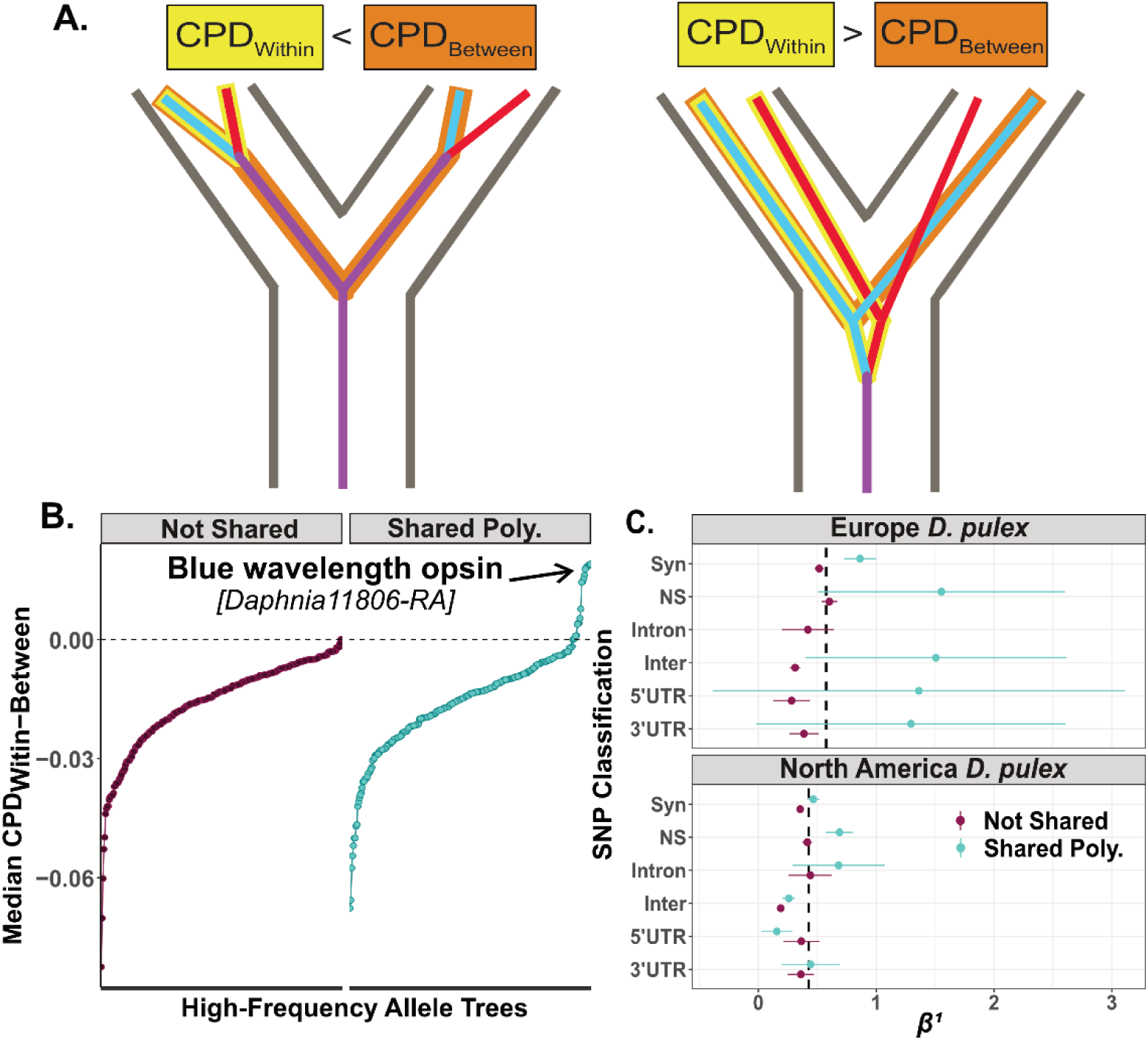
Convergent evolution, trans-specificity, and signatures of balancing selection. **A)** Visualization of two adaptive hypotheses that produce shared polymorphisms, convergent evolution on the left and trans-specificity on the right. For each tree, we calculated the median pairwise cophenetic distance as the distance within species (CPD_Within_; yellow highlighted pair) – between species (CPD_Between_; orange highlight) for shared polymorphisms and non-shared polymorphisms. CPD_Within-Between_ < 0 describes the consensus species-tree topology (Left), while CPD_Within-Between_ > 0 describes an allele-specific tree topology consistent with an old mutation being maintained within the sequence (Right). The red and blue branches indicate examples of shared polymorphisms between species. B) CPD_Within-Between_ for non-synonymous shared SNPs and non-shared SNPs above 0.25 minor allele frequency (MAF) in both species. Each allele-tree was made from 30 samples from North American and European *D. pulex*. At the focal SNP, we extracted 500bps surrounding the focal SNP. C) *β^1^* is a statistic that detects balancing selection. We show the mean with 95% standard errors for several SNP classifications (SYN=synonymous, NS=non-synonymous, Intron=intronic, Inter=intergenic, 5’ UTR=5’ untranslated region, 3’ UTR=3’ untranslated region). The dotted vertical line is the average *β^1^* within each species.

### Selective forces acting on shared polymorphisms

European and North American *D. pulex* possess an excess of shared polymorphism relative to neutral or demographic models, suggesting that some form of selection could be maintaining these polymorphisms. We sought to identify old-balanced polymorphisms and convergently evolved polymorphisms by building allele-trees surrounding focal shared polymorphisms. If shared polymorphisms arose via convergence, then allele-trees would be concordant with the species-tree and all parental haplotypes from the same species would be reciprocally monophyletic (Figure 4A). If shared polymorphisms arose prior to the species split, then allele trees will not necessarily be concordant with the species tree. Notably, if there are multiple shared polymorphisms in close linkage, then alleles from the two species will cluster together and be distinct from the species tree. However, it is important to note that if only a single trans-specific polymorphism is the target of balancing selection that arose prior to the species split, then recombination could have eroded the signal of linked ancient polymorphism and the allele trees will be concordant with the species tree (Gao et al., 2015). Thus, our analysis cannot accurately separate convergence from trans-specificity in all cases, but can identify genes that have multiple, linked shared trans-specific polymorphisms that could be the target of long-term balancing selection.

We summarized allele tree discordance by calculating the pairwise cophenetic distances (CPD, Cardona et al., 2013) within and between haplotypes of the same-species from allele-trees that contain high-frequency (MAF>0.25), non-synonymous, shared polymorphisms. When allele trees resemble the species-tree topology, the within-species distances will be lower than the between-species distances (CPD_Within_-CPD_Between_ < 0; Figure 4A). However, if alleles from two species cluster together, and are discordant with the species-tree, the within-species distances will be larger than between-species distances (CPD_Within_-CPD_Between_ > 0). A small number of allele trees surrounding shared polymorphisms have a positive CPD_Within-Between_ value, consistent with balancing selection maintaining trans-specific haplotypes (Figure 4A). However, most shared polymorphisms have negative CPD_Within-Between_ values (Figure 4B), consistent either with convergently evolution or trans-specificity. Although determining the fraction of shared polymorphisms that arose via either selective mechanism is challenging, it seems unlikely that all shared polymorphisms with negative CPD_Within-Between_ arose via convergent adaptive evolution. This is because the probability of mutation occurring at the same nucleotide in two species is small (μ^2^ ∼ 10^-17^; Keith et al., 2016), coupled with the low establishment probability for anything but the most strongly beneficial mutations.

Regardless of whether shared polymorphisms arose via convergent evolution, or prior to the species split, they could have been subject to balancing selection. To test this hypothesis, we first calculated *α_b_*, a statistic to estimate the proportion of non-synonymous sites under balancing selection using a contingency table odds ratio of both private-species’ alleles and shared polymorphisms (Soni et al., 2022). We found that *α_b_* is significantly positive across the genome, indicating that balancing selection is influencing non-synonymous shared polymorphisms (*α_b_*=0.082 [0.05, 0.114], *p*=1.5x10^-6^). Next, we calculated *β^1^*, a site-frequency spectrum-based statistic for detecting signals of balancing selection (Siewert & Voight, 2017) at both shared and control SNPs. We found that *β^1^* at shared polymorphisms are significantly higher than zero in both species for non-synonymous SNPs (one sample t-test: Euro. *t* = 7.8, *df* = 270, *p* = 1.75x10^-13^; NAm. *t* = 18.8, *df* = 1563, *p* = 2.2x10^-16^; Figure 4C). Shared synonymous sites are also significantly elevated *β^1^* in both species (NAm. *t* = 25.12, *df* = 4367, *p* = 2.2x10^-16^; Euro. *t* = 20.41, *df* = 1481, *p* = 2.2x10^-16^; Figure 4C).

### Trans-specific polymorphisms at a blue wavelength opsin affect behavior and show evidence of genetic overdominance

Of the common, non-synonymous shared polymorphisms, 14 (5%) have positive CPD_Within-Between_ values (Figure 4B). Almost all of these shared polymorphisms (13/14) are within a rhabdomeric blue wavelength opsin (BLOP) gene (Brandon et al., 2017). The BLOP that we identify is found as a single copy in European and North American *D. pulex* (Supplemental Figure 4A). The 13 non-synonymous shared SNPs reside across several exons (Figure 5A) and encompass a large linkage block within European *D. pulex* (*r^2^* > 0.7 ∼ 1.5kbps; Figure 5B&C), thereby explaining the allele-tree species-tree discordance (Figure 4B; Figure 5C) and suggest that these alleles are trans-specific polymorphisms (TSP) that predate the split between North American and European *D. pulex*.

**Figure 5.**
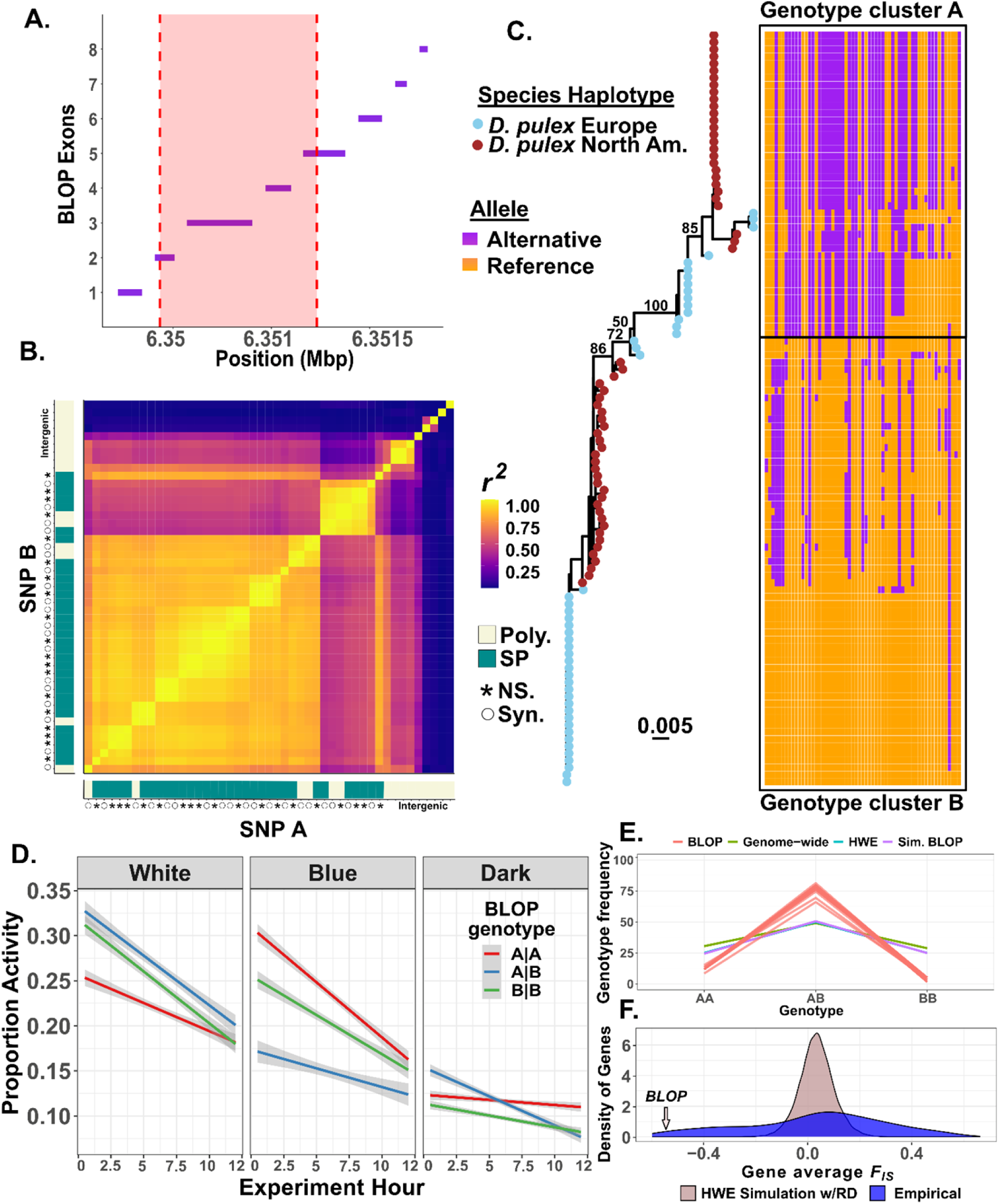
Behavioral and fitness effects of trans-specific SNPs at a blue-light opsin. **A)** Gene structure showing the length and position of exons within the BLOP (*Daphnia11806*). The shaded red region indicates the location of a large high-linkage block identified in panel B. **B)** Pairwise linkage disequilibrium (*r^2^*) for every SNP within the BLOP for European *D. pulex*, filtered for SNPs with a MAF > 0.01. The right and bottom tile objects indicate whether the SNP is polymorphic (Poly; khaki) or shared polymorphism (SP; blue-green). NS refers to non-synonymous polymorphism and Syn refers to synonymous polymorphism represented by asterisks and open circles respectively. **C)** Allele-tree made from the gene for a subset of phased samples of North American and European *D. pulex*. Tip symbols indicate whether the samples are North American or European *D. pulex*. Numbers indicate bootstrap support. The included haplotype plot and multiple-sequence alignment showcase the presence of each SNP within the gene, colored for whether the allele is derived (purple) or reference (gold). **D)** The activity of individual European *D. pulex* was measured for 12 hours for three genotypes in three different light conditions. Lines represent the best fit and 95% standard errors. **E)** Average segregation frequency of F1 genotypes expected based on a double heterozygous cross (i.e., AB x AB) using empirical read depth at each SNP. “BLOP” is the empirical segregation of trans-specific polymorphisms within the blue wavelength opsin gene among F1 genotypes. “Genome-wide” is the segregation for SNPs based on the read depth. “HWE” is the segregation pattern expected for Hardy Weinberg equilibrium. “Sim. BLOP” is the segregation pattern expected for the SNPs within the blue opsin gene based on empirical read depth. **F)** Distribution of average gene *F_IS_*. “HWE Simulation w/RD” is the expected *F_IS_* for each gene based on the empirical read depth for each SNP within every gene and “Empirical” is the average *F_IS_* across genes. The small arrow denotes where the gene average for the blue wavelength opsin falls along the empirical distribution.

If these haplotypes at the BLOP have been maintained since prior to the split between North American and European *D. pulex* 10 MYA (Figure 1C), they may have a functional effect. To test this hypothesis, we measured the light-induced activity of European *D. pulex* clones that harbor distinct haplotypes bearing alternate shared alleles. We first assigned clonal haplotypes to one of two genetic clusters (Figure 5C) and tested the activity levels of all three genotypes (AA, AB, BB) in different light conditions. We found that genotype has a significant effect on activity that is dependent on light conditions (*χ^2^*=5,849.71, *df*=4, *p* < 2x10^-16^; Supplemental Table 3). In general, all genotypes had low activity in dark conditions. Heterozygotes have the highest activity levels when exposed to white light yet have the lowest activity when exposed to blue light consistent with shifts between genetic overdominance and underdominance affecting behavior (Figure 5D).

Overdominance affecting behavior could also translate into overdominance affecting fitness. If trans-specific polymorphisms at the BLOP cause overdominance in fitness, heterozygotes should be more common than expected under Hardy-Weinberg equilibrium. To test the hypothesis, we examined segregation patterns of trans-specific SNPs at the BLOP among F1 offspring derived from a cross between two clones that are both heterozygous for the trans-specific haplotypes we identified. These clones were previously referred to as “super-clone A” and “super-clone C” by Barnard-Kubow et al., (2022). Both clones had reached high frequency in the southern English (Dorset) pond D8 by the end of the 2017 growing season. In 2018, most individuals in the D8 pond were the F1 offspring between super-clone A and C enabling us to directly test if there is an excess of heterozygotes relative to the expected Mendelian segregation patterns among the F1s. First, we calculated the frequency of AA, AB, and BB genotypes at trans-specific polymorphisms, without downsampling to one clone per MLG, at the BLOP. We find that there is a strong excess of heterozygotes in the wild-caught individuals compared to expectations from Hardy-Weinberg (HWE) and compared to random SNPs in the genome or other TSPs (Figure 5E). Next, we calculated the distribution of *F_IS_*, a measure of the departure of HWE, at genes across the genome and found that the BLOP gene is amongst the most strongly negative *F_IS_* compared to other genes (*F_IS_* = -0.54; Figure 5E). Indeed, the BLOP has amongst the smallest 2.6% of *F_IS_* values that we measured. Even if we examine the genotype distribution by only sampling one genotype per clonal lineage we still observe an excess of heterozygotes (Supplemental Figure 5C&F), again suggesting natural selection in the wild. We also examined genotype frequencies in lab-generated AxC and CxC F1s. In contrast to our field-sampled individuals, we do not observe an excess of heterozygotes from a lab-generated cross of the same clones (Supplemental Figure 5A-E).

## Discussion

In this study, we examined the evolutionary forces that generate and maintain shared polymorphisms in the *D. pulex* species complex. This species complex contains several taxa that have played a preeminent role in evolutionary genetics and ecology, yet their phylogenetic relationship and nomenclature has proven challenging for over 150 years. Here, we used whole-genome sequences coupled with polymorphism data to resolve the nuclear phylogeny of members of this species group, to evaluate mechanisms that can generate shared polymorphisms between species, and to test the functional and fitness effects of ancient mutations. We show that there is an excess of shared polymorphisms between North American and European *D. pulex* that cannot be explained by neutral or demographic processes, thereby implicating some form of natural selection as a force maintaining polymorphism. For one gene, a blue wavelength opsin, we show that shared polymorphism is likely ancient, predating speciation, and has functional consequences on behavior and fitness in the wild.

### Phylogenetics of the D. pulex species group

Members of the genus *Daphnia*, and the *D. pulex* species group in particular, have proven challenging from a taxonomic perspective since their early description. For instance, Leydig separated *D. pulex* from *D. magna* and *D. longispina* (Leydig, 1860, p. 117), but did not further describe divisions in the group. Richard (1896) identified *D. obtusa* as a distinct species from *D. pulex* (p. 260), but also described ten subspecies of *D. pulex* found across the Americas and Eurasia (p. 232-255). Scourfield (1942) reinforced the view that *D. obtusa* and *D. pulex* are distinct species and emphasized the view that this species group represents several lineages in various stages of speciation. Johnson (1951), in his description of British members of the *D. pulex* group, noted that American forms resembling species in the *D. pulex* group are not likely monophyletic with Eurasian species of the same name, although these naming conventions have persisted (Brooks, 1957b; Omilian & Lynch, 2009; Ye et al., 2023). The challenge of morphological classification in the *D. pulex* group stems from a limited number of diagnostic characteristics (Brooks, 1957b; Dodson, 1981), coupled with phenotypic plasticity (Colbourne et al., 1997), mating type variation (Heier & Dudycha, 2009; Jose & Dufresne, 2010), and cytological variation (Gómez et al., 2016; Hosseinie, 1966). However, recent phylogenetic analysis of mitochondrial markers has shown the *D. pulex* group consists of many distinct lineages and that the deepest splits within the *D. pulex* species group occur between Eurasian and North American taxa (Crease et al., 2012; Ye et al., 2022). Consistent with these results, allopatric speciation has been estimated to account for roughly 40% of cladogenetic events within *Daphnia* (Adamowicz et al., 2009), a process possibly enhanced by cycles of glaciation (Chin & Cristescu, 2021). We show that substantial genetic division exists between North American and European taxa and that these taxa are separated by millions of years (Figure 1). Given the relatively deep split time between members of the *D. pulex* species group, it is likely that they have distinct features ranging from their response to environmental stimuli to their impact on the ecosystem. Further study of the behavioral, physiological, and ecological interactions of these taxa is warranted.

The complicated nature of the *D. pulex* species group is compounded by incomplete reproductive isolation between them. North American *D. pulex* and North American *D. pulicaria* are known to hybridize in the wild (Xu et al., 2015; Ye et al., 2019). Hybrids between these lineages are obligately asexual and fail to produce functional males (Tucker et al., 2013; Xu et al., 2015; Ye et al., 2019). These post-zygotic reproductive incompatibilities are a hallmark of taxa undergoing incipient speciation (Coughlan & Matute, 2020). Consistent with this view, we show that the split time between North American *D. pulex* and North American *D. pulicaria* based on the nuclear genome is recent, within 3 million years (Figure 1C). Our estimate is consistent with a study made from mitochondrial genomes (Colbourne et al., 1998), but older than another using a limited number of nuclear markers (Omilian & Lynch, 2009). Nonetheless, genomic data clearly show that hybridization between these North American lineages occurs (Figure 2). Previous analysis of mitochondrial markers placed European *D. pulicaria* as sister to the North American *D. pulex/pulicaria* clade (e.g., Marková et al., 2013), a result consistent with the nuclear phylogeny we constructed (Figure 1C). European *D. pulicaria* also shows evidence of hybridization with members of the North American *D. pulex/pulicaria* clade (Figure 2B&E), although such hybridization is not likely recent or could have occurred with other lineages in this complex. Although the North American taxa, along with European *D. pulicaria* show signals of hybridization with each other, European *D. pulex* appears to be a well-defined species. We show that European *D. pulex* split from the other *D. pulex/pulicaria* taxa approximately 10 million years ago (Figure 1C) and has little to no evidence of recent hybridization (Figure 2A&E).

### The generation of shared polymorphisms

Polymorphisms that are shared between species represent a particularly interesting class of mutation because they can reflect a wide variety of evolutionary processes. On the one hand, shared polymorphisms could reflect neutral processes when they occur between closely related species. For example, species that diverged relatively recently will share many polymorphisms because of incomplete lineage sorting (Hobolth et al., 2011) or ongoing gene-flow (Payseur & Rieseberg, 2016). While the presence of neutral shared polymorphisms due to incomplete lineage sorting or gene-flow is important for understanding features such as historical population size (Suh et al., 2015) or barriers to migration (Kutschera et al., 2014), they can obscure selective forces such as convergent adaptive evolution or balancing selection that can also generate or maintain shared polymorphism. Therefore, to examine these selective forces, it is important to identify species that have diverged long enough ago that incomplete lineage sorting and ongoing gene-flow are limited. Our work identifies European and North American *D. pulex* as two such species because of their deep split time and limited evidence for hybridization.

We show that there are tens of thousands of polymorphisms that are shared between European and North American *D. pulex* (Figure 3A, Supplemental Table 2) and suggest that natural selection is responsible for their presence. Natural selection has often been implicated as playing a key role in maintaining shared polymorphism. For instance, polymorphisms at MHC genes in vertebrates are routinely identified to be older than the species split (Aguilar et al., 2004; Azevedo et al., 2015; Klein et al., 1993) and are thought to be maintained as polymorphism via mechanisms such as negative frequency dependence or genetic overdominance (Key et al., 2014). In other cases, shared polymorphisms in a variety of taxa have possibly arisen via convergent evolution to common selective pressures such as pathogens (Těšický & Vinkler, 2015) and have been maintained in both species via balancing selection (Solberg et al., 2008). North American and European *D. pulex* genes involved in the immune system do not show any systematic evidence of shared polymorphism (results not shown), although the ratio of non-synonymous to synonymous polymorphisms is higher for shared polymorphisms (0.58) than non-shared polymorphisms in either North American *D. pulex* or European *D. pulex* (0.53 and 0.46, respectively; see Supplemental Table 2). Therefore, it is likely that many of these shared polymorphisms are functional and subject to some form of balancing selection.

The shared non-synonymous polymorphisms that we identified have allele trees that largely reflect the species tree (Figure 4A). Taken at face value, this result is consistent with convergent evolution. Others have suggested that widespread convergent evolution is an unlikely mechanism generating shared polymorphisms (Klein et al., 1993). Is this conclusion valid for *Daphnia*? The probability of a beneficial mutation arising in a population is a function of its census size (Pennings & Hermisson, 2006) and its establishment in a population is a function of the selective value of the mutation (Haldane, 1927). While the long-term effective population size of both European and North American *D. pulex* is somewhat limited (N_e_ < 1 million; Supplemental Figure 3), the census size at any single pond or lake can be quite large, possibly reaching into the millions of individuals (Dudycha, 2004), while the global census size of either species can reach upwards of 10^12^ individuals (Buffalo, 2021). Therefore, across the species range, these taxa are not likely mutation-limited. Indeed, recurrent *de novo* evolution of beneficial mutations have been hypothesized to occur rapidly and contribute to within-population variation in male production rates (Barnard-Kubow et al., 2022) and morphological responses to predators (Becker et al., 2022). Temporally and spatially variable natural selection have also been shown to be a potent force acting on *Daphnia* populations (Chaturvedi et al., 2021; Lynch, 1987; Lynch et al., 2023), suggesting that positive selection on new beneficial mutations could be strong enough to prevent beneficial mutations from being lost (Flynn et al., 2017). Therefore, it is conceivable that such shared polymorphisms between North American and European *D. pulex* arose independently. On the other hand, distinguishing between convergent evolution and old trans-specific polymorphism based on comparisons between allele trees and species trees is not always possible. This is especially so when only a single trans-specific polymorphism is the direct target of selection. In this scenario, the linked neutral trans-specific polymorphisms that generate the footprint of genealogical discordance will be eroded via recombination. Regardless of whether the many shared polymorphisms that we observe between North American and European *D. pulex* arose via convergent evolution or have been maintained since prior to the species split, these mutations tend to be associated with signatures of elevated polymorphism (Figure 4C), suggestive of balancing selection, as seen in other systems (Leffler et al., 2013).

### Natural selection maintains functional trans-specific polymorphisms in a blue wavelength opsin gene

We show that one gene, a blue wavelength opsin harbors trans-specific mutations that predates the split between North American and European *D. pulex* (Figure 4B, 5C). At this locus, allele trees differ from species trees, a signal that is consistent with trans-specific polymorphism (Charlesworth, 2006; Fijarczyk & Babik, 2015). This BLOP gene has 15 non-synonymous TSPs and extensive heterozygosity (Supplemental Figure 4B). The extensive heterozygosity and linkage structure of this BLOP makes it a high priority candidate for functional characterization. Research into the North American *D. pulex* genome has shown ancient expansion of opsin genes in general that occurred over 145 mya (Brandon et al., 2017). Recent work showcases that positive selection strength is distinct between North American *D. pulex* and *D. pulicaria* at opsin genes highlighting the complex patterns of selection acting upon opsins across the genome (Ye et al., 2023). It could be that this blue wavelength opsin mediates behavioral responses like predator avoidance or vertical diel migration seen in most Daphnids (Li et al., 2022). Our laboratory experimental work shows that alternate genotypes at the BLOP have different behavioral activity patterns in response to different light conditions (Figure 5D). Indeed, it even appears that there are changes in dominance as a function of light treatment, a feature that is consistent with the long-term persistence of balanced polymorphisms (Wittmann et al., 2017).

Our genomic analyses show that there is an excess of heterozygotes at this locus. Likewise, our experimental work identified a putative fitness advantage in the wild (Figure 5E). These results are consistent with previous experiments and observations in *Daphnia* (Haag & Ebert, 2007; Hebert et al., 1982). Our result relies on temporal sampling of a single wild population along with the reconstruction of the pedigree using genomic data of wild-caught individuals (Barnard-Kubow et al., 2022). Barnard-Kubow *et al.,* (2022) show that two clones became dominant in a pond and then crossed with each other, producing a population of F1 offspring the following year. The two dominant clones were heterozygous for the trans-specific SNPs at the BLOP and thus we expect their offspring to follow a simple Mendelian 1:2:1 ratio. In contrast, we observe an excess of heterozygous individuals in the population. This pattern is largely explained by heterozygous clones reaching higher frequency in the population by the time they were sampled suggesting that heterozygotes had higher fitness and thus were more likely to survive. By contrasting genotype frequencies from the field to the lab (Supplemental Figure 5), we conclude that the excess of heterozygotes in the field is not likely due to factors such as inbreeding depression or associative overdominance (Ohta, 1971). Instead, these patterns likely emerged due to the action of natural selection. Given the strong link between looming stimulus, movement, and predator avoidance in *Daphnia* (Pijanowska & Kowalczewski, 1997; Ringelberg, 1999; Van Gool & Ringelberg, 2003), we hypothesize that trans-specific polymorphisms at the BLOP locus may play a role in conferring a fitness advantage by reducing encounters with predators or by facilitating migration through the water column.

## Conclusion

Our study elucidates the evolutionary history and genetic structure of the *D. pulex* species complex and provides evidence that shared polymorphisms are common between cryptic species. We show that balancing selection broadly influences shared polymorphisms and that a small fraction predates the species-split. We experimentally study the functional significance of shared polymorphisms across specific ecological contexts and show that these polymorphisms are associated with fitness in the wild. While we present four hypotheses related to the origin and maintenance of shared polymorphism (hybridization, incomplete lineage sorting, convergence, and balancing selection), these hypotheses are not mutually exclusive. Additionally, the evolutionary mechanisms presented as hypotheses will all be affected by background levels of recombination, historic shifts in *N_e_*, and patterns of positive and purifying selection acting upon the genome (Charlesworth, 2009; Charlesworth, 2006). Despite this challenge, we laid the groundwork for understanding the mechanisms by which genetic diversity is maintained between cryptic *D. pulex* species.

## Author contributions

CSM: Conceptualization, Data curation, Formal analysis, Investigation, Methodology, Project administration, Software, Visualization, Writing - original draft, Writing - review & editing. MK: Methodology, Investigation, Writing - review & editing. DJB: Formal analysis, Software, Visualization, Writing - review & editing. MD: Investigation, Writing - review & editing. DB: Investigation, Resources, Writing - review & editing. JCBN: Formal analysis, Software, Methodology, Writing - review & editing. AR: Formal analysis, Software, Writing - review & editing. AOB: Conceptualization, Formal analysis, Funding acquisition, Investigation, Methodology, Project administration, Software, Supervision, Validation, Visualization, Writing - review & editing

## Acknowledgments

The authors acknowledge members of the Bergland lab for their feedback related to the manuscript’s development. We thank Benedict Adam Lenhart, Daniel Nondorf, Christopher Robinson, Zoë Ogilvie, and Kendall Branham for their valuable comments on early versions of the manuscript. We also thank Dieter Ebert for his advice about the divergence-time estimates. The authors acknowledge Research Computing at UVA for providing computational resources and technical support that have contributed to the results reported within this manuscript. URL: https://rc.virginia.edu.

---

**Supplemental Figure 1.**
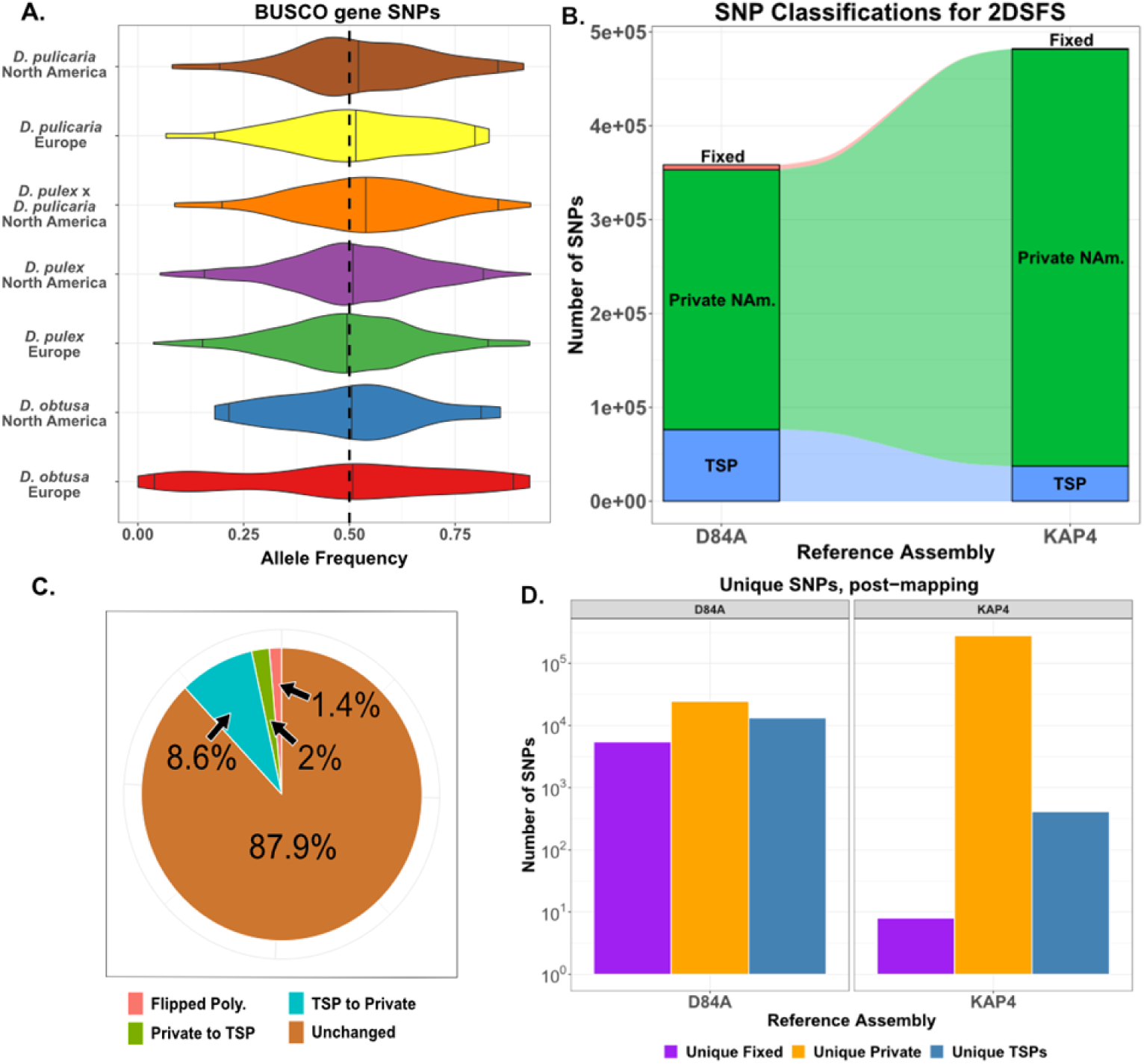
No evidence of reference allele bias across the *Daphnia pulex* species complex. **A)** For each species, we extracted representative individuals (ranging from n=5-100 depending on the number of samples per species) and 1,000 biallelic heterozygous BUSCO gene SNPs (100 bootstraps) to gauge the severity of reference allele bias across the genome. We calculated the proportion of the alternative and reference dosage within a given individual for each site. The x-axis measures the proportion of alternative to reference dosage for each SNP and we show the 95% quantiles and median. **B)** Alluvial plot of the SNP classifications between assemblies of the European *D. pulex* (D84A) and the North American *D. pulex* (KAP4). **C)** Proportion of SNP classification changes when mapping to KAP4 exclusively. **D)** The number of classified SNPs that are exclusive to each assembly.

**Supplemental Figure 2.**
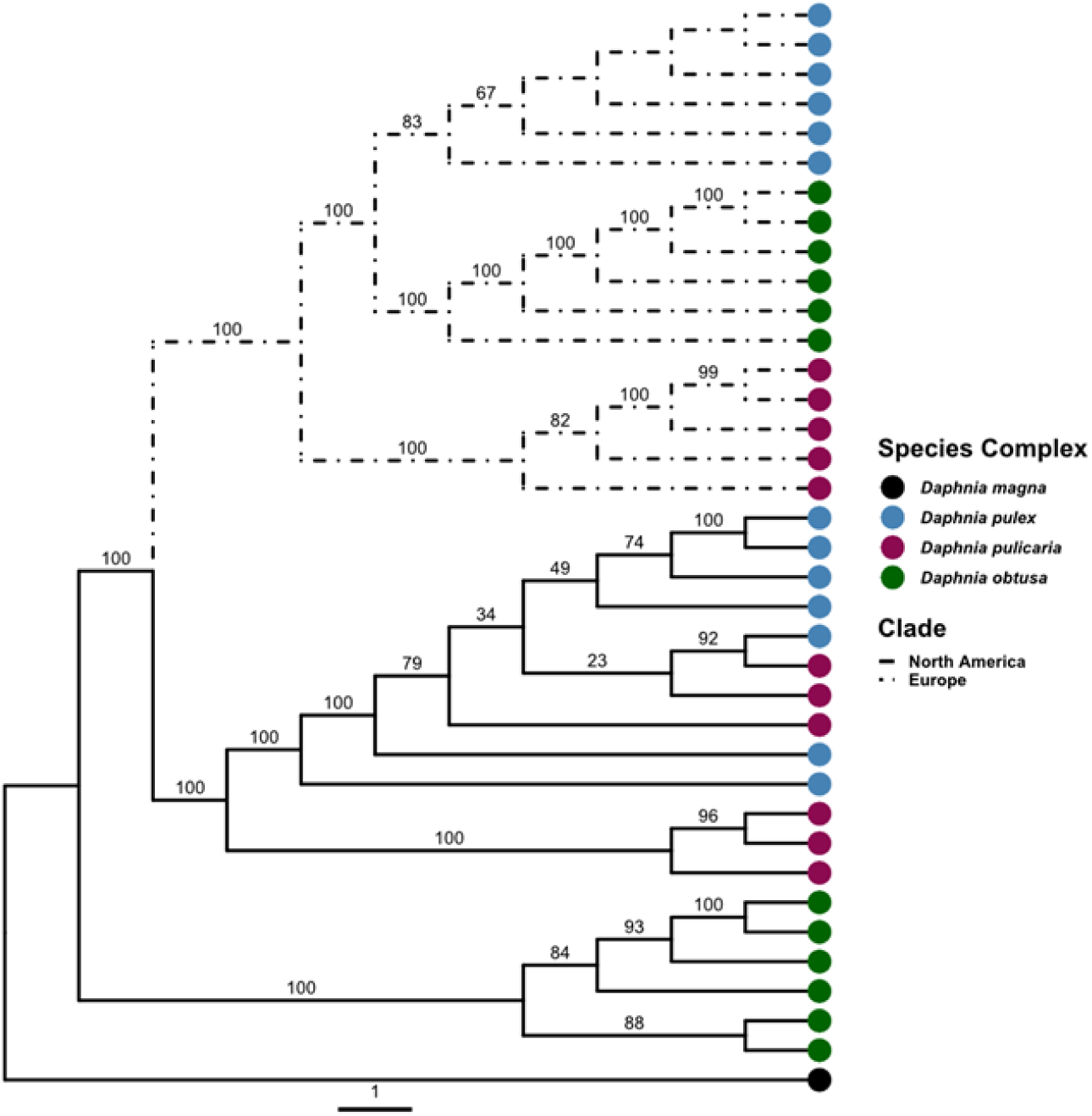
Mitochondrial protein-coding tree for the *D. pulex* species complex. A maximum-likelihood tree with the “TN+F+I+G4” model output from IQTree2. The tree is rooted with *D. magna* as an outgroup. Bootstrap supports are listed as node labels.

**Supplemental Figure 3.**
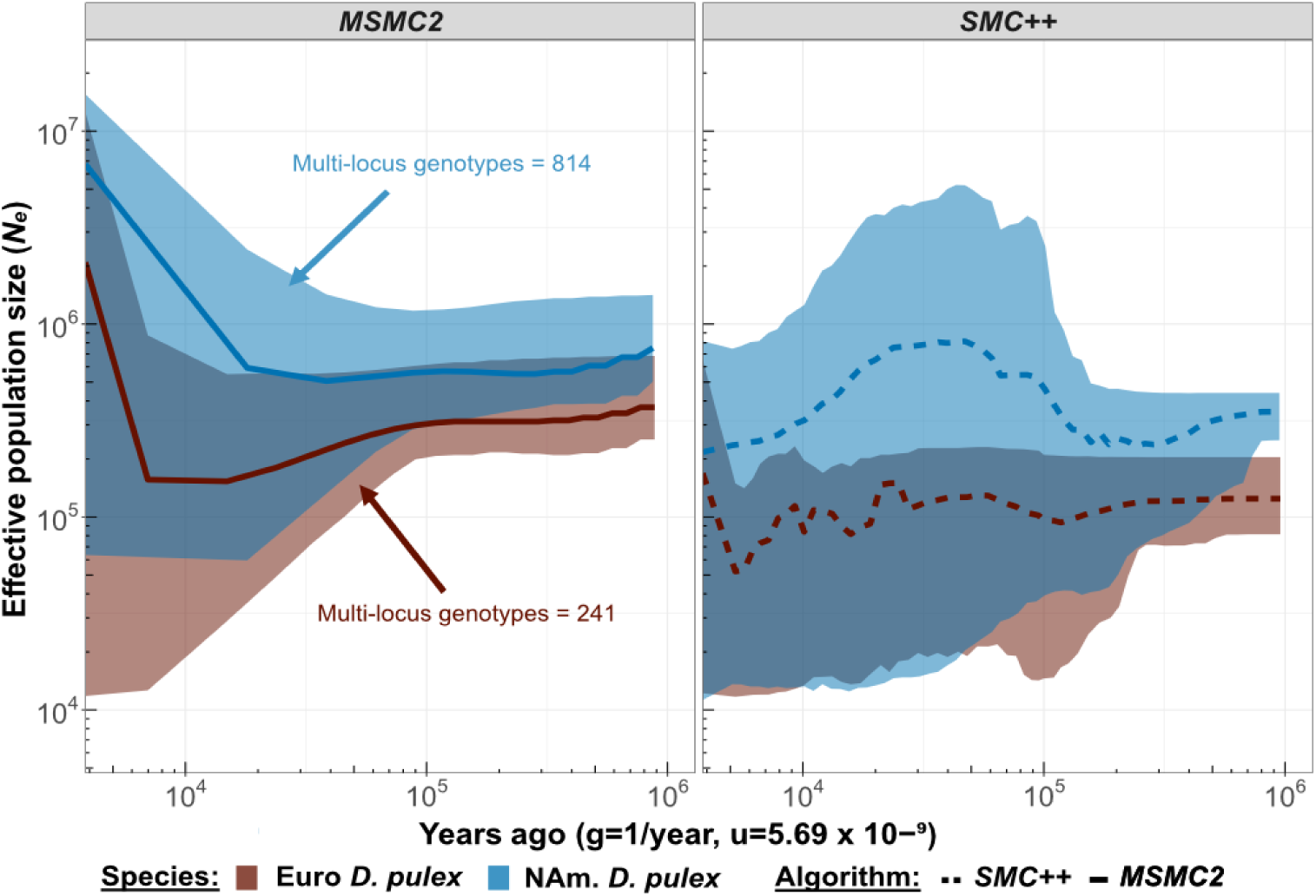
Demographic reconstruction of North American and European *D. pulex* species. MSMC2 and SMC++ output for each multi-locus genotype sample. Each multi-locus genotype sample was run independently. The shaded ribbon shows the upper 95% quantiles and lower 5% quantiles from the run estimates.

**Supplemental Figure 4.**
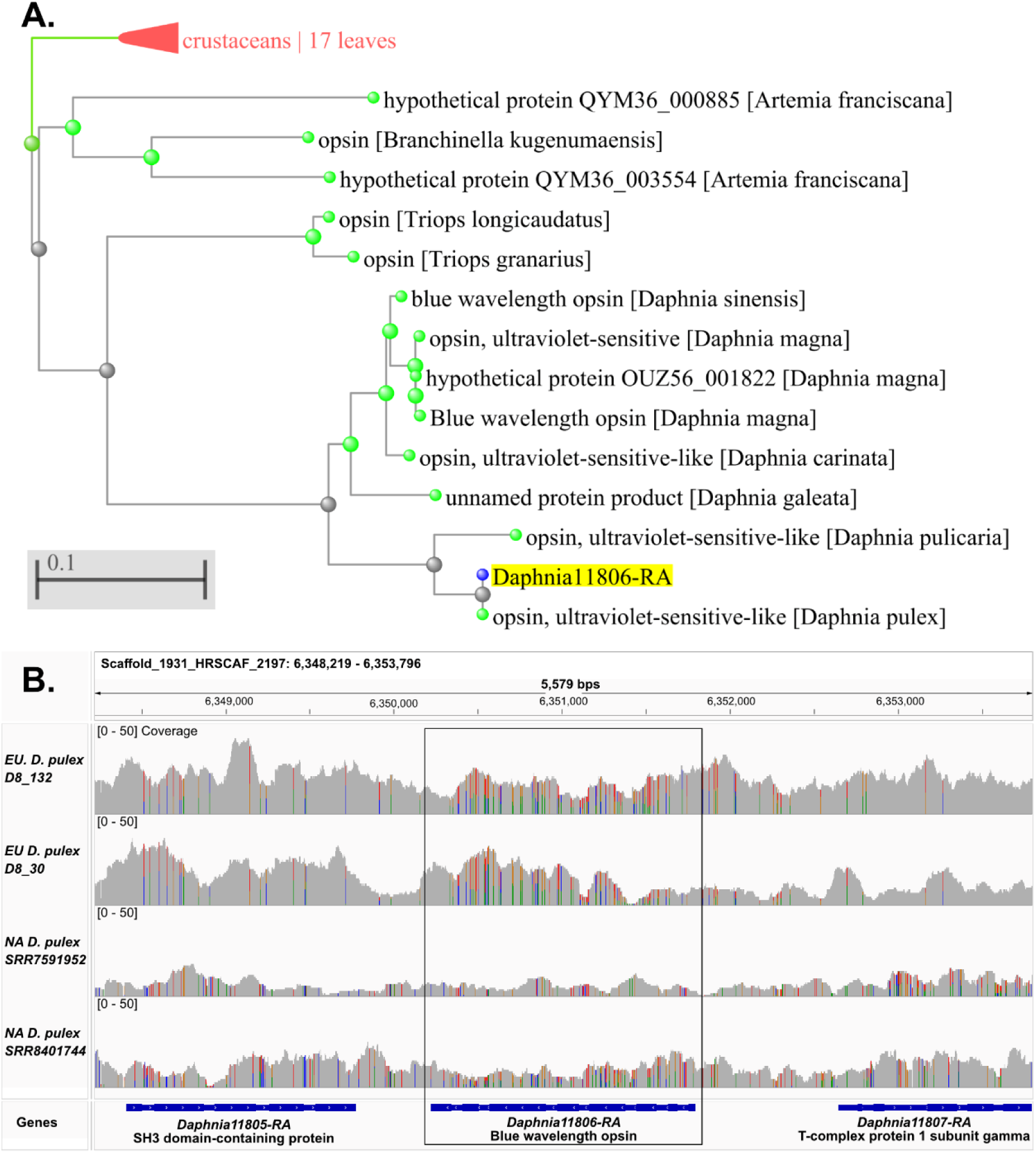
Blue wavelength opsin gene and orthologous proteins in Crustacea and within-species heterozygosity. **A)** This neighbor-joining protein tree was generated using *Blast’s* tree widget. The query sequence is highlighted in yellow and has a blue tip symbol. The green tip symbols are related Crustacean protein sequences with the species name in brackets. **B)** We subsampled two representative individuals within the European (EU) and North American (NA) *D. pulex* species and are showing the coverage for each individual set to [0-50]. The vertical-colored bars are heterozygous regions (i.e., split-colored bars) and homozygous alternative alleles (i.e., whole-colored bars), gray base pairs are the reference allele.

**Supplemental Figure 5.**
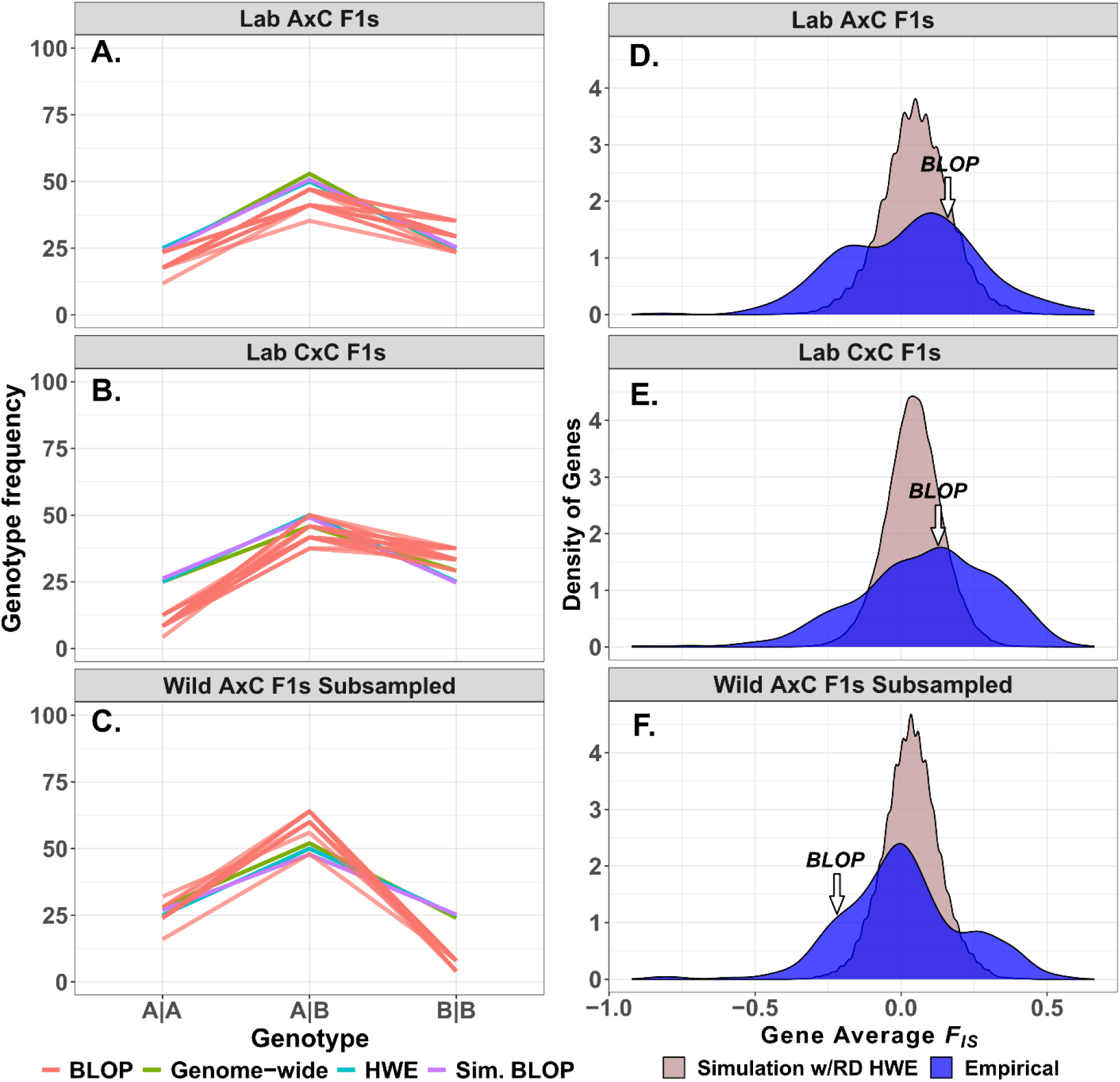
Segregation patterns and *F_IS_* of polymorphisms across lab and wild crossed *Daphnia* clones. **A-C)** Average segregation frequency of F1 genotypes expected based on a double heterozygous cross (i.e., AB x AB) using empirical read depth at each SNP. We produced crosses of AxC in the lab shown in panel A and CxC shown in panel B. Panel C shows the F1 genotypes subsampled based on their status belonging to superclones identified in Barnard-Kubow et al. 2022, reflecting a conservative sampling approach. “Genome-wide” is the segregation for SNPs based on the read depth. “HWE” is the segregation pattern expected for Hardy Weinberg equilibrium. “Sim. BLOP” is the segregation pattern expected for the SNPs within the blue opsin gene based on empirical read depth. “BLOP” is the empirical segregation of trans-specific polymorphisms within the blue wavelength opsin gene among F1 genotypes. **D-F)** Distribution of average gene *F_IS_*. “HWE Simulation w/RD” is the expected *F_IS_* for each gene based on the empirical read depth for each SNP within every gene and “Empirical” is the average is the *F_IS_*across genes. The small arrow denotes where the gene average for the blue wavelength opsin falls along the empirical distribution.

## References

1. Adamowicz, S. J., Petrusek, A., Colbourne, J. K., Hebert, P. D. N., & Witt, J. D. S. (2009). The scale of divergence: A phylogenetic appraisal of intercontinental allopatric speciation in a passively dispersed freshwater zooplankton genus. Molecular Phylogenetics and Evolution, 50(3), 423–436. 10.1016/j.ympev.2008.11.026

2. Aguilar, A., Roemer, G., Debenham, S., Binns, M., Garcelon, D., & Wayne, R. K. (2004). High MHC diversity maintained by balancing selection in an otherwise genetically monomorphic mammal. Proceedings of the National Academy of Sciences, 101(10), 3490–3494. 10.1073/pnas.0306582101

3. Alexander, D. H., & Lange, K. (2011). Enhancements to the ADMIXTURE algorithm for individual ancestry estimation. BMC Bioinformatics, 12. 10.1186/1471-2105-12-246

4. Andrews, S. (2010). FastQC: A Quality Control Tool for High Throughput Sequence Data. Available online at: http://www.bioinformatics.babraham.ac.uk/projects/fastqc/

5. Azevedo, L., Serrano, C., Amorim, A., & Cooper, D. N. (2015). Trans-species polymorphism in humans and the great apes is generally maintained by balancing selection that modulates the host immune response. Human Genomics, 9(1), 21. 10.1186/s40246-015-0043-1

6. Barnard-Kubow, K. B., Becker, D., Murray, C. S., Porter, R., Gutierrez, G., Erickson, P., Nunez, J. C. B., Voss, E., Suryamohan, K., Ratan, A., Beckerman, A., & Bergland, A. O. (2022). Genetic Variation in Reproductive Investment Across an Ephemerality Gradient in *Daphnia pulex*. Molecular Biology and Evolution, 39(6), msac121. 10.1093/molbev/msac121

7. Bates, D., Mächler, M., Bolker, B., & Walker, S. (2015). Fitting Linear Mixed-Effects Models Using lme4. Journal of Statistical Software, 67(1). 10.18637/jss.v067.i01

8. Baym, M., Kryazhimskiy, S., Lieberman, T. D., Chung, H., Desai, M. M., & Kishony, R. (2015). Inexpensive Multiplexed Library Preparation for Megabase-Sized Genomes. PLOS ONE, 10(5), Article 5. 10.1371/journal.pone.0128036

9. Becker, D., Barnard-Kubow, K., Porter, R., Edwards, A., Voss, E., Beckerman, A. P., & Bergland, A. O. (2022). Adaptive phenotypic plasticity is under stabilizing selection in *Daphnia*. Nature Ecology & Evolution, 6(10), 1449–1457. 10.1038/s41559-022-01837-5

10. Bergland, A. O., Behrman, E. L., O’Brien, K. R., Schmidt, P. S., & Petrov, D. A. (2014). Genomic Evidence of Rapid and Stable Adaptive Oscillations over Seasonal Time Scales in *Drosophila*. PLoS Genetics, 10(11), e1004775. 10.1371/journal.pgen.1004775

11. Bernt, M., Donath, A., Jühling, F., Externbrink, F., Florentz, C., Fritzsch, G., Pütz, J., Middendorf, M., & Stadler, P. F. (2013). MITOS: Improved de novo metazoan mitochondrial genome annotation. Molecular Phylogenetics and Evolution, 69(2), 313–319. 10.1016/j.ympev.2012.08.023

12. Bolger, A. M., Lohse, M., & Usadel, B. (2014). Trimmomatic: A flexible trimmer for Illumina sequence data. Bioinformatics, 30(15), 2114–2120. 10.1093/bioinformatics/btu170

13. Bouckaert, R., Heled, J., Kühnert, D., Vaughan, T., Wu, C. H., Xie, D., Suchard, M. A., Rambaut, A., & Drummond, A. J. (2014). BEAST 2: A Software Platform for Bayesian Evolutionary Analysis. PLoS Computational Biology, 10(4), 1–6. 10.1371/journal.pcbi.1003537

14. Brandon, C. S., Greenwold, M. J., & Dudycha, J. L. (2017). Ancient and Recent Duplications Support Functional Diversity of *Daphnia* Opsins. Journal of Molecular Evolution, 84(1), 12–28. 10.1007/s00239-016-9777-1

15. Brooks, J. L. (1957a). The species problem in freshwater animals. Pp. 81–123, in: The Species Problem. Amer. Assoc. Advan. Sci.

16. Brooks, J. L. (1957b). The systematics of North American *Daphnia*. Memoirs of the Connecticut Academy of Arts & Science.

17. Buffalo, V. (2021). Quantifying the relationship between genetic diversity and population size suggests natural selection cannot explain Lewontin’s Paradox. eLife, 10, e67509. 10.7554/eLife.67509

18. Cardona, G., Mir, A., Rosselló, F., Rotger, L., & Sánchez, D. (2013). Cophenetic metrics for phylogenetic trees, after Sokal and Rohlf. BMC Bioinformatics, 14(1), 3. 10.1186/1471-2105-14-3

19. Castoe, T. A., De Koning, A. P. J., Kim, H.-M., Gu, W., Noonan, B. P., Naylor, G., Jiang, Z. J., Parkinson, C. L., & Pollock, D. D. (2009). Evidence for an ancient adaptive episode of convergent molecular evolution. Proceedings of the National Academy of Sciences, 106(22), 8986–8991. 10.1073/pnas.0900233106

20. Černý, M., & Hebert, P. D. N. (1999). Intercontinental allozyme differentiation among four holarctic *Daphnia* species. Limnology and Oceanography, 44(6), 1381–1387. 10.4319/lo.1999.44.6.1381

21. Charlesworth, B. (2009). Effective population size and patterns of molecular evolution and variation. Nature Reviews Genetics, 10(3), 195–205. 10.1038/nrg2526

22. Charlesworth, D. (2006). Balancing selection and its effects on sequences in nearby genome regions. PLoS Genetics, 2(4), 379–384. 10.1371/journal.pgen.0020064

23. Chaturvedi, A., Zhou, J., Raeymaekers, J. A. M., Czypionka, T., Orsini, L., Jackson, C. E., Spanier, K. I., Shaw, J. R., Colbourne, J. K., & De Meester, L. (2021). Extensive standing genetic variation from a small number of founders enables rapid adaptation in *Daphnia*. Nature Communications, 12(1), 4306. 10.1038/s41467-021-24581-z

24. Chen, N.-C., Solomon, B., Mun, T., Iyer, S., & Langmead, B. (2021). Reference flow: Reducing reference bias using multiple population genomes. Genome Biology, 22(1), 8. 10.1186/s13059-020-02229-3

25. Chin, T. A., & Cristescu, M. E. (2021). Speciation in *Daphnia*. Molecular Ecology, mec.15824. 10.1111/mec.15824

26. Chiu, J. C., Low, K. H., Pike, D. H., Yildirim, E., & Edery, I. (2010). Assaying Locomotor Activity to Study Circadian Rhythms and Sleep Parameters in *Drosophila*. Journal of Visualized Experiments, 43, 2157. 10.3791/2157

27. Cingolani, P., Platts, A., Wang, L. L., Coon, M., Nguyen, T., Wang, L., Land, S. J., Lu, X., & Ruden, D. M. (2012). A program for annotating and predicting the effects of single nucleotide polymorphisms, SnpEff: SNPs in the genome of Drosophila melanogaster strain w^1118^ ; iso-2; iso-3. Fly, 6(2), 80–92. 10.4161/fly.19695

28. Clark, A. G. (1997). Neutral behavior of shared polymorphism. Proceedings of the National Academy of Sciences, 94(15), 7730–7734. 10.1073/pnas.94.15.7730

29. Colbourne, J. K., Crease, T. J., Weider, L. J., Hebert, P. D. N., Duferesne, F., & Hobaek, A. (1998). Phylogenetics and evolution of a circumarctic species complex (Cladocera: *Daphnia pulex*). Biological Journal of the Linnean Society, 65(3), 347–365. 10.1111/j.1095-8312.1998.tb01146.x

30. Colbourne, J. K., Pfrender, M. E., Gilbert, D., Thomas, W. K., Tucker, A., Oakley, T. H., Tokishita, S., Aerts, A., Arnold, G. J., Basu, M. K., Bauer, D. J., Caceres, C. E., Carmel, L., Casola, C., Choi, J.-H., Detter, J. C., Dong, Q., Dusheyko, S., Eads, B. D., … Boore, J. L. (2011). The Ecoresponsive Genome of *Daphnia pulex*. Science, 331(6017), 555–561. 10.1126/science.1197761

31. Colbourne, J. K.J, Hebert, P. D., Taylor, D. J., & Givnish, T. J. (1997). Evolutionary origins of phenotypic diversity in *Daphnia*. Molecular evolution and adaptive radiation, 163–188.

32. Cornetti, L., Fields, P. D., Van Damme, K., & Ebert, D. (2019). A fossil-calibrated phylogenomic analysis of *Daphnia* and the Daphniidae. Molecular Phylogenetics and Evolution, 137, 250–262. 10.1016/j.ympev.2019.05.018

33. Coughlan, J. M., & Matute, D. R. (2020). The importance of intrinsic postzygotic barriers throughout the speciation process. Philosophical Transactions of the Royal Society B: Biological Sciences, 375(1806), 20190533. 10.1098/rstb.2019.0533

34. Crease, T. J., Omilian, A. R., Costanzo, K. S., & Taylor, D. J. (2012). Transcontinental Phylogeography of the *Daphnia pulex* Species Complex. PLoS ONE, 7(10), e46620. 10.1371/journal.pone.0046620

35. Crow, J. F. & Kimura, M. (1970). An introduction to population genetics theory. New York, NY: Harper & Row, Publishers, Inc.

36. Daniel, F., Revolution Analytics, & Weston, S. (2022). doMC: Foreach Parallel Adaptor for “parallel.”

37. Decaestecker, E., Gaba, S., Raeymaekers, J. A. M., Stoks, R., Van Kerckhoven, L., Ebert, D., & De Meester, L. (2007). Host–parasite ‘Red Queen’ dynamics archived in pond sediment. Nature, 450(7171), 870–873. 10.1038/nature06291

38. Dodson, S. I. (1981). Morphological variation of *Daphnia pulex* Leydig (Crustacea: Cladocera) and related species from North America. Hydrobiologia, 83(1), 101–114. 10.1007/BF02187155

39. Dowle, M., & Srinivasan, A. (2023). data.table: Extension of “data.frame.” Https://R-Datatable.Com, Https://Rdatatable.Gitlab.Io/Data.Table, Https://Github.Com/Rdatatable/Data.Table.

40. Dudycha, J. L. (2004). Mortality dynamics of *Daphnia* in contrasting habitats and their role in ecological divergence. Freshwater Biology, 49(5), 505–514. 10.1111/j.1365-2427.2004.01201.x

41. Ebert, D. (2022). *Daphnia* as a versatile model system in ecology and evolution. EvoDevo, 13(1), 16. 10.1186/s13227-022-00199-0

42. Edwards, ScottV., & Beerli, P. (2000). Perspective: Gene Divergence, Population Divergence, and the Variance in Coalescence Time in Phylogeographic Studies. Evolution, 54(6), 1839–1854. 10.1111/j.0014-3820.2000.tb01231.x

43. Emms, D. M., & Kelly, S. (2019). OrthoFinder: Phylogenetic orthology inference for comparative genomics. Genome Biology, 20(1), 238. 10.1186/s13059-019-1832-y

44. Erickson, P. A., Weller, C. A., Song, D. Y., Bangerter, A. S., Schmidt, P., & Bergland, A. O. (2020). Unique genetic signatures of local adaptation over space and time for diapause, an ecologically relevant complex trait, in *Drosophila melanogaster*. PLOS Genetics, 16(11), e1009110. 10.1371/journal.pgen.1009110

45. Ewels, P., Magnusson, M., Lundin, S., & Käller, M. (2016). MultiQC: Summarize analysis results for multiple tools and samples in a single report. Bioinformatics, 32(19), 3047–3048. 10.1093/bioinformatics/btw354

46. Flynn, J. M., Chain, F. J. J., Schoen, D. J., & Cristescu, M. E. (2017). Spontaneous Mutation Accumulation in *Daphnia pulex* in Selection-Free vs. Competitive Environments. Molecular Biology and Evolution, 34(1), 160–173. 10.1093/molbev/msw234

47. Gao, Z., Przeworski, M., & Sella, G. (2015). Footprints of ancient-balanced polymorphisms in genetic variation data from closely related species. Evolution, 69(2), 431–446. 10.1111/evo.12567

48. Garnier, S., Ross, N., BoB Rudis, Filipovic-Pierucci, A., Galili, T., Timelyportfolio, Greenwell, B., Sievert, C., Harris, D. J., & JJ Chen. (2021). *sjmgarnier/viridis: Viridis 0.6.0 (pre-CRAN release)* (v0.6.0pre). 10.5281/ZENODO.4679424

49. Geoffrey Fryer. (1991). Functional morphology and the adaptive radiation of the Daphniidae (Branchiopoda: Anomopoda). Philosophical Transactions of the Royal Society of London. Series B: Biological Sciences, 331(1259), 1–99. 10.1098/rstb.1991.0001

50. Gómez, R., Van Damme, K., Gosálvez, J., Morán, E. S., & Colbourne, J. K. (2016). Male meiosis in Crustacea: Synapsis, recombination, epigenetics and fertility in *Daphnia magna*. Chromosoma, 125(4), 769–787. 10.1007/s00412-015-0558-1

51. Günther, T., & Nettelblad, C. (2019). The presence and impact of reference bias on population genomic studies of prehistoric human populations. PLOS Genetics, 15(7), e1008302. 10.1371/journal.pgen.1008302

52. Haag, C. R., & Ebert, D. (2007). Genotypic selection in *Daphnia* populations consisting of inbred sibships. Journal of Evolutionary Biology, 20(3), 881–891. 10.1111/j.1420-9101.2007.01313.x

53. Haldane, J. B. S. (1927). A Mathematical Theory of Natural and Artificial Selection, Part V: Selection and Mutation. Mathematical Proceedings of the Cambridge Philosophical Society, 23(7), 838–844. 10.1017/S0305004100015644

54. Harris, RS. (2007). Improved Pairwise Alignment of Genomic DNA. The Pennsylvania State University, University Park, PA.

55. Hebert, P. D. N., Ferrari, D. C., & Crease, T. J. (1982). Heterosis in *Daphnia*: A Reassessment. The American Naturalist, 119(3), 427–434. 10.1086/283921

56. Hebert, P. D. N., & Wilson, C. C. (1994). Provincialism in plankton: endemism and allopatric speciation in australian *Daphnia*. Evolution, 48(4), 1333–1349. 10.1111/j.1558-5646.1994.tb05317.x

57. Hedrick, P. W. (2013). Adaptive introgression in animals: Examples and comparison to new mutation and standing variation as sources of adaptive variation. Molecular Ecology, 22(18), 4606–4618. 10.1111/mec.12415

58. Heier, C. R., & Dudycha, J. L. (2009). Ecological speciation in a cyclic parthenogen: Sexual capability of experimental hybrids between *Daphnia pulex* and *Daphnia pulicaria*. Limnology and Oceanography, 54(2), 492–502. 10.4319/lo.2009.54.2.0492

59. Held, C., Koenemann, S., & Schubart, C. D. (Eds.). (2016). Phylogeography and Population Genetics in Crustacea (0 ed.). CRC Press. 10.1201/b11113

60. Hoang, D. T., Chernomor, O., von Haeseler, A., Minh, B. Q., & Vinh, L. S. (2018). UFBoot2: Improving the Ultrafast Bootstrap Approximation. Molecular Biology and Evolution, 35(2), 518–522. 10.1093/molbev/msx281

61. Hobolth, A., Dutheil, J. Y., Hawks, J., Schierup, M. H., & Mailund, T. (2011). Incomplete lineage sorting patterns among human, chimpanzee, and orangutan suggest recent orangutan speciation and widespread selection. Genome Research, 21(3), 349–356. 10.1101/gr.114751.110

62. Hosseinie, Farammarz. (1966). The ecology and reproductive cytology of *Daphnia middendorffiana*, FischerCladocera) from the arctic. Indiana University ProQuest Dissertations Publishing.

63. Huerta-Sánchez, E., Jin, X., Asan, Bianba, Z., Peter, B. M., Vinckenbosch, N., Liang, Y., Yi, X., He, M., Somel, M., Ni, P., Wang, B., Ou, X., Huasang, Luosang, J., Cuo, Z. X. P., Li, K., Gao, G., Yin, Y., … Nielsen, R. (2014). Altitude adaptation in Tibetans caused by introgression of Denisovan-like DNA. Nature, 512(7513), 194–197. 10.1038/nature13408

64. Jackson, C. E., Xu, S., Ye, Z., Pfrender, M. E., Lynch, M., Colbourne, J. K., & Shaw, J. R. (2021). Chromosomal rearrangements preserve adaptive divergence in ecological speciation. BioRxiv, 2021–08. 10.1101/2021.08.20.457158

65. Johnson, D. S. (1951). A study of the physiology and ecology of certain Cladocera. University of London, Bedford College (United Kingdom) ProQuest Dissertations Publishing.

66. Jose, C., & Dufresne, F. (2010). Differential survival among genotypes of *Daphnia pulex* differing in reproductive mode, ploidy level, and geographic origin. Evolutionary Ecology, 24(2), 413–421. 10.1007/s10682-009-9314-4

67. Jouganous, J., Long, W., Ragsdale, A. P., & Gravel, S. (2017). Inferring the Joint Demographic History of Multiple Populations: Beyond the Diffusion Approximation. Genetics, 206(3), 1549–1567. 10.1534/genetics.117.200493

68. Kalyaanamoorthy, S., Minh, B. Q., Wong, T. K. F., von Haeseler, A., & Jermiin, L. S. (2017). ModelFinder: Fast model selection for accurate phylogenetic estimates. Nature Methods, 14(6), 587–589. 10.1038/nmeth.4285

69. Kamvar, Z. N., Tabima, J. F., & Grünwald, N. J. (2014). *Poppr*: An R package for genetic analysis of populations with clonal, partially clonal, and/or sexual reproduction. PeerJ, 2, e281. 10.7717/peerj.281

70. Katoh, K., & Standley, D. M. (2013). MAFFT Multiple Sequence Alignment Software Version 7: Improvements in Performance and Usability. Molecular Biology and Evolution, 30(4), 772–780. 10.1093/molbev/mst010

71. Keith, N., Tucker, A. E., Jackson, C. E., Sung, W., Lucas Lledó, J. I., Schrider, D. R., Schaack, S., Dudycha, J. L., Ackerman, M., Younge, A. J., Shaw, J. R., & Lynch, M. (2016). High mutational rates of large-scale duplication and deletion in *Daphnia pulex*. Genome Research, 26(1), 60–69. 10.1101/gr.191338.115

72. Key, F. M., Teixeira, J. C., de Filippo, C., & Andrés, A. M. (2014). Advantageous diversity maintained by balancing selection in humans. Current Opinion in Genetics & Development, 29, 45–51. 10.1016/j.gde.2014.08.001

73. Klein, J., Sato, A., Nagl, S., & O’hUigín, C. (1998). Molecular trans-species polymorphism. Annual Review of Ecology and Systematics, 29(1), 1–21. 10.1146/annurev.ecolsys.29.1.1

74. Klein, J., Satta, Y., Takahata, N., & O’hUigin, C. (1993). Trans-specific *Mhc* polymorphism and the origin of species in primates. Journal of Medical Primatology, 22(1), 57–64. 10.1111/j.1600-0684.1993.tb00637.x

75. Koenig, D., Hagmann, J., Li, R., Bemm, F., Slotte, T., Neuffer, B., Wright, S. I., & Weigel, D. (2019). Long-term balancing selection drives evolution of immunity genes in Capsella. eLife, 8, e43606. 10.7554/eLife.43606

76. Kutschera, V. E., Bidon, T., Hailer, F., Rodi, J. L., Fain, S. R., & Janke, A. (2014). Bears in a Forest of Gene Trees: Phylogenetic Inference Is Complicated by Incomplete Lineage Sorting and Gene Flow. Molecular Biology and Evolution, 31(8), 2004–2017. 10.1093/molbev/msu186

77. Lee, B. T., Barber, G. P., Benet-Pagès, A., Casper, J., Clawson, H., Diekhans, M., Fischer, C., Gonzalez, J. N., Hinrichs, A. S., Lee, C. M., Muthuraman, P., Nassar, L. R., Nguy, B., Pereira, T., Perez, G., Raney, B. J., Rosenbloom, K. R., Schmelter, D., Speir, M. L., … Kent, W. J. (2022). The UCSC Genome Browser database: 2022 update. Nucleic Acids Research, 50(D1), D1115–D1122. 10.1093/nar/gkab959

78. Leffler, E. M., Gao, Z., Pfeifer, S., Segurel, L., Auton, A., Venn, O., Bowden, R., Bontrop, R., Wall, J. D., Sella, G., Donnelly, P., McVean, G., & Przeworski, M. (2013). Multiple Instances of Ancient Balancing Selection Shared Between Humans and Chimpanzees. Science, 339(6127), 1578–1582. 10.1126/science.1234070

79. Leinonen, R., Sugawara, H., Shumway, M., & on behalf of the International Nucleotide Sequence Database Collaboration. (2011). The Sequence Read Archive. Nucleic Acids Research, 39(Database), D19–D21. 10.1093/nar/gkq1019

80. Leydig, F. (1860). Naturgeschichte der Daphniden:(Crustacea cladocera). Laupp & Siebeck.

81. Li, D., Huang, J., Zhou, Q., Gu, L., Sun, Y., Zhang, L., & Yang, Z. (2022). Artificial Light Pollution with Different Wavelengths at Night Interferes with Development, Reproduction, and Antipredator Defenses of *Daphnia magna*. Environmental Science & Technology, 56(3), 1702–1712. 10.1021/acs.est.1c06286

82. Li, H. (2013). Aligning sequence reads, clone sequences and assembly contigs with BWA-MEM. arXiv preprint arXiv:1303.3997

83. Li, H., Handsaker, B., Wysoker, A., Fennell, T., Ruan, J., Homer, N., Marth, G., Abecasis, G., Durbin, R., & 1000 Genome Project Data Processing Subgroup. (2009). The Sequence Alignment/Map format and SAMtools. Bioinformatics, 25(16), 2078–2079. 10.1093/bioinformatics/btp352

84. Lynch, M. (1987). The Consequences of Fluctuating Selection for Isozyme Polymorphisms in *Daphnia*. Genetics, 115(4), 657–669. 10.1093/genetics/115.4.657

85. Lynch, M., Gutenkunst, R., Ackerman, M., Spitze, K., Ye, Z., Maruki, T., & Jia, Z. (2017). Population Genomics of *Daphnia pulex*. Genetics, 206(1), 315–332. 10.1534/genetics.116.190611

86. Lynch, M., Wei, W., Ye, Z., & Pfrender, M. (2023). The Genome-wide Signature of Short-term Temporal Selection. bioRxiv, 2023.04.28.538790. 10.1101/2023.04.28.538790

87. Malinsky, M., Matschiner, M., & Svardal, H. (2021). Dsuite—Fast D-statistics and related admixture evidence from VCF files. Molecular Ecology Resources, 21(2), 584–595. 10.1111/1755-0998.13265

88. Marková, S., Dufresne, F., Manca, M., & Kotlík, P. (2013). Mitochondrial Capture Misleads about Ecological Speciation in the *Daphnia pulex* Complex. PLOS ONE, 8(7), e69497. 10.1371/journal.pone.0069497

89. Martin, M., Ebert, P., & Marschall, T. (2023). Read-Based Phasing and Analysis of Phased Variants with WhatsHap. In B. A. Peters & R. Drmanac (Eds.), Haplotyping (Vol. 2590, pp. 127–138). Springer US. 10.1007/978-1-0716-2819-5_8

90. Mayer, W. E., Jonker, M., Klein, D., Ivanyi, P., Van Seventer, G., & Klein, J. (1988). Nucleotide sequences of chimpanzee MHC class I alleles: Evidence for trans-species mode of evolution. The EMBO Journal, 7(9), 2765–2774. 10.1002/j.1460-2075.1988.tb03131.x

91. McCoy, R. C., Garud, N. R., Kelley, J. L., Boggs, C. L., & Petrov, D. A. (2014). Genomic inference accurately predicts the timing and severity of a recent bottleneck in a nonmodel insect population. Molecular Ecology, 23(1), 136–150. 10.1111/mec.12591

92. Mi, H., Muruganujan, A., Casagrande, J. T., & Thomas, P. D. (2013). Large-scale gene function analysis with the PANTHER classification system. Nature Protocols, 8(8), 1551–1566. 10.1038/nprot.2013.092

93. Novikova, P. Y., Hohmann, N., Nizhynska, V., Tsuchimatsu, T., Ali, J., Muir, G., Guggisberg, A., Paape, T., Schmid, K., Fedorenko, O. M., Holm, S., Säll, T., Schlötterer, C., Marhold, K., Widmer, A., Sese, J., Shimizu, K. K., Weigel, D., Krämer, U., … Nordborg, M. (2016). Sequencing of the genus *Arabidopsis* identifies a complex history of nonbifurcating speciation and abundant trans-specific polymorphism. Nature Genetics, 48(9), 1077–1082. 10.1038/ng.3617

94. Nunez, J. C. B., Rong, S., Damian-Serrano, A., Burley, J. T., Elyanow, R. G., Ferranti, D. A., Neil, K. B., Glenner, H., Rosenblad, M. A., Blomberg, A., Johannesson, K., & Rand, D. M. (2021). Ecological Load and Balancing Selection in Circumboreal Barnacles. Molecular Biology and Evolution, 38(2), 676–685. 10.1093/molbev/msaa227

95. Ohta, T. (1971). Associative overdominance caused by linked detrimental mutations. Genetical Research, 18(3), 277–286. 10.1017/S0016672300012684

96. Omilian, A. R., & Lynch, M. (2009). Patterns of Intraspecific DNA Variation in the *Daphnia* Nuclear Genome. Genetics, 182(1), 325–336. 10.1534/genetics.108.099549

97. Pantel, J. H., Juenger, T. E., & Leibold, M. A. (2011). Environmental gradients structure *Daphnia pulex* × *pulicaria* clonal distribution: Hybrid *Daphnia* clonal distribution. Journal of Evolutionary Biology, 24(4), 723–732. 10.1111/j.1420-9101.2010.02196.x

98. Paradis, E., & Schliep, K. (2019). ape 5.0: An environment for modern phylogenetics and evolutionary analyses in R. Bioinformatics, 35(3), 526–528. 10.1093/bioinformatics/bty633

99. Payseur, B. A., & Rieseberg, L. H. (2016). A genomic perspective on hybridization and speciation. Molecular Ecology, 25(11), 2337–2360. 10.1111/mec.13557

100. Pennings, P. S., & Hermisson, J. (2006). Soft Sweeps II—Molecular Population Genetics of Adaptation from Recurrent Mutation or Migration. Molecular Biology and Evolution, 23(5), 1076–1084. 10.1093/molbev/msj117

101. Pijanowska, J., & Kowalczewski, A. (1997). Predators can induce swarming behaviour and locomotory responses in *Daphnia*. Freshwater Biology, 37(3), 649–656. 10.1046/j.1365-2427.1997.00192.x

102. Poplin, R., Ruano-Rubio, V., DePristo, M. A., Fennell, T. J., Carneiro, M. O., Van Der Auwera, G. A., Kling, D. E., Gauthier, L. D., Levy-Moonshine, A., Roazen, D., Shakir, K., Thibault, J., Chandran, S., Whelan, C., Lek, M., Gabriel, S., Daly, M. J., Neale, B., MacArthur, D. G., & Banks, E. (2017). Scaling accurate genetic variant discovery to tens of thousands of samples. BioRxiv, 201178. 10.1101/201178

103. R Core Development Team. (2013). R: A language and environment for statistical computing.

104. Rambaut, A., Drummond, A. J., Xie, D., Baele, G., & Suchard, M. A. (2018). Posterior Summarization in Bayesian Phylogenetics Using Tracer 1.7. Systematic Biology, 67(5), 901–904. 10.1093/sysbio/syy032

105. Revell, L. J. (2019). *learnPopGen*: An R package for population genetic simulation and numerical analysis. Ecology and Evolution, 9(14), 7896–7902. 10.1002/ece3.5412

106. Richard J. (1896). Révision des Cladocères. Deuxième Partie. Anomopoda. Famille III.— Daphnidae (Vol. 2). Ann. Sci. Nat. Zool.

107. Ringelberg, J. (1999). The photobehaviour of *Daphnia spp*. As a model to explain diel vertical migration in zooplankton. Biological Reviews of the Cambridge Philosophical Society, 74(4), 397–423. 10.1017/S0006323199005381

108. Sayers, E. W., Bolton, E. E., Brister, J. R., Canese, K., Chan, J., Comeau, D. C., Connor, R., Funk, K., Kelly, C., Kim, S., Madej, T., Marchler-Bauer, A., Lanczycki, C., Lathrop, S., Lu, Z., Thibaud-Nissen, F., Murphy, T., Phan, L., Skripchenko, Y., … Sherry, S. T. (2022). Database resources of the national center for biotechnology information. Nucleic Acids Research, 50(D1), D20–D26. 10.1093/nar/gkab1112

109. Schiffels, S., & Wang, K. (2020). MSMC and MSMC2: The Multiple Sequentially Markovian Coalescent. In J. Y. Dutheil (Ed.), Statistical Population Genomics (Vol. 2090, pp. 147–166). Springer US. 10.1007/978-1-0716-0199-0_7

110. Scourfield, D. J. (1942). XIX.—The “Pulex” forms of *Daphnia* and their Separation into Two Distinct Series. Annals and Magazine of Natural History, 9(51), 202–219. 10.1080/03745481.1942.9755477

111. Ségurel, L., Gao, Z., & Przeworski, M. (2013). Ancestry runs deeper than blood: The evolutionary history of *ABO* points to cryptic variation of functional importance. BioEssays, 35(10), 862–867. 10.1002/bies.201300030

112. Ségurel, L., Thompson, E. E., Flutre, T., Lovstad, J., Venkat, A., Margulis, S. W., Moyse, J., Ross, S., Gamble, K., Sella, G., Ober, C., & Przeworski, M. (2012). The ABO blood group is a trans-species polymorphism in primates. Proceedings of the National Academy of Sciences, 109(45), 18493–18498. 10.1073/pnas.1210603109

113. Seppey, M., Manni, M., & Zdobnov, E. M. (2019). BUSCO: Assessing Genome Assembly and Annotation Completeness. In M. Kollmar (Ed.), Gene Prediction (Vol. 1962, pp. 227–245). Springer New York. 10.1007/978-1-4939-9173-0_14

114. Shen, W., Le, S., Li, Y., & Hu, F. (2016). SeqKit: A Cross-Platform and Ultrafast Toolkit for FASTA/Q File Manipulation. PLOS ONE, 11(10), e0163962. 10.1371/journal.pone.0163962

115. Siewert, K. M., & Voight, B. F. (2017). Detecting Long-Term Balancing Selection Using Allele Frequency Correlation. Molecular Biology and Evolution, 34(11), 2996–3005. 10.1093/molbev/msx209

116. Simão, F. A., Waterhouse, R. M., Ioannidis, P., Kriventseva, E. V., & Zdobnov, E. M. (2015). BUSCO: Assessing genome assembly and annotation completeness with single-copy orthologs. Bioinformatics, 31(19), 3210–3212. 10.1093/bioinformatics/btv351

117. Slater, G., & Birney, E. (2005). Automated generation of heuristics for biological sequence comparison. BMC Bioinformatics, 6(1), 31. 10.1186/1471-2105-6-31

118. So, M., Ohtsuki, H., Makino, W., Ishida, S., Kumagai, H., Yamaki, K. G., & Urabe, J. (2015). Invasion and molecular evolution of *Daphnia pulex* in Japan. Limnology and Oceanography, 60(4), 1129–1138. 10.1002/lno.10087

119. Solberg, O. D., Mack, S. J., Lancaster, A. K., Single, R. M., Tsai, Y., Sanchez-Mazas, A., & Thomson, G. (2008). Balancing selection and heterogeneity across the classical human leukocyte antigen loci: A meta-analytic review of 497 population studies. Human Immunology, 69(7), 443–464. 10.1016/j.humimm.2008.05.001

120. Soni, V., Vos, M., & Eyre-Walker, A. (2022). A new test suggests hundreds of amino acid polymorphisms in humans are subject to balancing selection. PLOS Biology, 20(6), e3001645. 10.1371/journal.pbio.3001645

121. Spitze, K. (1993). Population structure in *Daphnia obtusa*: Quantitative genetic and allozymic variation. Genetics, 135(2), 367–374. 10.1093/genetics/135.2.367

122. Standard, A. (2007). Standard guide for conducting acute toxicity tests on test materials with fishes, macroinvertebrates, and amphibians. West Conshohocken, PA, United States. DOI: 0.1520/E0729-96. URL: Www. Atsm. Org.

123. Suh, A., Smeds, L., & Ellegren, H. (2015). The Dynamics of Incomplete Lineage Sorting across the Ancient Adaptive Radiation of Neoavian Birds. PLOS Biology, 13(8), e1002224. 10.1371/journal.pbio.1002224

124. Tarailo-Graovac, M., & Chen, N. (2009). Using RepeatMasker to Identify Repetitive Elements in Genomic Sequences. Current Protocols in Bioinformatics, 25(1). 10.1002/0471250953.bi0410s25

125. Terhorst, J., Kamm, J. A., & Song, Y. S. (2017). Robust and scalable inference of population history from hundreds of unphased whole genomes. Nature Genetics, 49(2), 303–309. 10.1038/ng.3748

126. Těšický, M., & Vinkler, M. (2015). Trans-Species Polymorphism in Immune Genes: General Pattern or MHC-Restricted Phenomenon? Journal of Immunology Research, 2015, 1–10. 10.1155/2015/838035

127. Thomas Lin Pedersen. (2022). patchwork: The Composer of Plots. Https://Patchwork.Data-Imaginist.Com, Https://Github.Com/Thomasp85/Patchwork.

128. Tucker, A. E., Ackerman, M. S., Eads, B. D., Xu, S., & Lynch, M. (2013). Population-genomic insights into the evolutionary origin and fate of obligately asexual *Daphnia pulex*. Proceedings of the National Academy of Sciences, 110(39), 15740–15745. 10.1073/pnas.1313388110

129. Unckless, R. L., Howick, V. M., & Lazzaro, B. P. (2016). Convergent Balancing Selection on an Antimicrobial Peptide in *Drosophila*. Current Biology, 26(2), 257–262. 10.1016/j.cub.2015.11.063

130. Van Gool, E., & Ringelberg, J. (2003). What goes down must come up: Symmetry in light-induced migration behaviour of *Daphnia*. Hydrobiologia, 491(1–3), 301–307. 10.1023/A:1024406324317

131. Villanueva, R. A. M., & Chen, Z. J. (2019). ggplot2: Elegant Graphics for Data Analysis (2nd ed.). Measurement: Interdisciplinary Research and Perspectives, 17(3), 160–167. 10.1080/15366367.2019.1565254

132. Wang, B., & Mitchell-Olds, T. (2017). Balancing selection and trans-specific polymorphisms. Genome Biology, 18(1), 231. 10.1186/s13059-017-1365-1

133. Wang, M., Zhang, L., Zhang, Z., Li, M., Wang, D., Zhang, X., Xi, Z., Keefover-Ring, K., Smart, L. B., DiFazio, S. P., Olson, M. S., Yin, T., Liu, J., & Ma, T. (2020). Phylogenomics of the genus *Populus* reveals extensive interspecific gene flow and balancing selection. New Phytologist, 225(3), 1370–1382. 10.1111/nph.16215

134. Wickham, H., Averick, M., Bryan, J., Chang, W., McGowan, L., François, R., Grolemund, G., Hayes, A., Henry, L., Hester, J., Kuhn, M., Pedersen, T., Miller, E., Bache, S., Müller, K., Ooms, J., Robinson, D., Seidel, D., Spinu, V., … Yutani, H. (2019). Welcome to the Tidyverse. Journal of Open Source Software, 4(43), 1686. 10.21105/joss.01686

135. Wills, C. (1975). Marginal overdominance in *Drosophila*. Genetics, 81(1), 177–189. 10.1093/genetics/81.1.177

136. Wittmann, M. J., Bergland, A. O., Feldman, M. W., Schmidt, P. S., & Petrov, D. A. (2017). Seasonally fluctuating selection can maintain polymorphism at many loci via segregation lift. Proceedings of the National Academy of Sciences, 114(46), E9932–E9941. 10.1073/pnas.1702994114

137. Wiuf, C., Zhao, K., Innan, H., & Nordborg, M. (2004). The Probability and Chromosomal Extent of trans-specific Polymorphism. Genetics, 168(4), 2363–2372. 10.1534/genetics.104.029488

138. Wu, Q., Han, T.-S., Chen, X., Chen, J.-F., Zou, Y.-P., Li, Z.-W., Xu, Y.-C., & Guo, Y.-L. (2017). Long-term balancing selection contributes to adaptation in *Arabidopsis* and its relatives. Genome Biology, 18(1), 217. 10.1186/s13059-017-1342-8

139. Xu, S., Li, L., Luo, X., Chen, M., Tang, W., Zhan, L., Dai, Z., Lam, T. T., Guan, Y., & Yu, G. (2022). *Ggtree*: A serialized data object for visualization of a phylogenetic tree and annotation data. iMeta, 1(4). 10.1002/imt2.56

140. Xu, S., Spitze, K., Ackerman, M. S., Ye, Z., Bright, L., Keith, N., Jackson, C. E., Shaw, J. R., & Lynch, M. (2015). Hybridization and the Origin of Contagious Asexuality in *Daphnia pulex*. *Molecular Biology and Evolution*, msv190. 10.1093/molbev/msv190

141. Ye, Z., Molinier, C., Zhao, C., Haag, C. R., & Lynch, M. (2019). Genetic control of male production in *Daphnia pulex*. Proceedings of the National Academy of Sciences, 116(31), 15602–15609. 10.1073/pnas.1903553116

142. Ye, Z., Pfrender, M. E., & Lynch, M. (2023a). Evolutionary Genomics of Sister Species Differing in Effective Population Sizes and Recombination Rates. Genome Biology and Evolution, 15(11), evad202. 10.1093/gbe/evad202

143. Ye, Z., Zhao, C., Raborn, R. T., Lin, M., Wei, W., Hao, Y., & Lynch, M. (2022). Genetic Diversity, Heteroplasmy, and Recombination in Mitochondrial Genomes of *Daphnia pulex* , *Daphnia pulicaria* , and *Daphnia obtusa*. Molecular Biology and Evolution, 39(4), msac059. 10.1093/molbev/msac059

144. Zhang, J., Kobert, K., Flouri, T., & Stamatakis, A. (2014). PEAR: A fast and accurate Illumina Paired-End reAd mergeR. Bioinformatics, 30(5), 614–620. 10.1093/bioinformatics/btt593

145. Zheng, X., Gogarten, S. M., Lawrence, M., Stilp, A., Conomos, M. P., Weir, B. S., Laurie, C., & Levine, D. (2017). SeqArray—A storage-efficient high-performance data format for WGS variant calls. Bioinformatics, 33(15), 2251–2257. 10.1093/bioinformatics/btx145

146. Zheng, X., Levine, D., Shen, J., Gogarten, S. M., Laurie, C., & Weir, B. S. (2012). A high-performance computing toolset for relatedness and principal component analysis of SNP data. Bioinformatics, 28(24), 3326–3328. 10.1093/bioinformatics/bts606

